# MDC1 counteracts restrained replication fork restart and its loss causes chemoresistance in BRCA1/2-deficient mammary tumors

**DOI:** 10.1101/2022.08.18.504391

**Authors:** Martin Liptay, Joana S. Barbosa, Ewa Gogola, Alexandra A. Duarte, Diego Dibitetto, Jonas A. Schmid, Ismar Klebic, Merve Mutlu, Myriam Siffert, Paola Francica, Israel Salguero, Marieke van de Ven, Renske de Korte-Grimmerink, Stephen P. Jackson, Jos Jonkers, Massimo Lopes, Sven Rottenberg

## Abstract

MDC1 is a key protein in DNA damage signaling. When DNA double-strand breaks (DSBs) occur, MDC1 localizes to sites of damage to promote the recruitment of other factors, including the 53BP1-mediated DSB repair pathway. By studying mechanisms of poly(ADP-ribose) polymerase inhibitor (PARPi) resistance in BRCA2;p53-deficient mouse mammary tumors, we identified a thus far unknown role of MDC1 in replication fork biology. MDC1 localizes at active replication forks during normal fork replication and its loss reduces fork speed. We show that MDC1 contributes to the restart of replication forks and thereby promotes sensitivity to PARPi and cisplatin. Loss of MDC1 causes MRE11-mediated resection, resulting in delayed fork restart. This improves DNA damage tolerance and causes chemoresistance in BRCA1/2-deficient cells. Hence, our results show a role for MDC1 in replication fork progression that mediates PARPi- and cisplatin-induced DNA damage, in addition to its role in DSB repair.

## INTRODUCTION

Despite the success of poly(ADP-ribose) polymerase (PARP) inhibitors in treatment of patients with BRCA1- or BRCA2-deficient tumors (Bryant et al., 2005; Farmer et al., 2005; Mateo et al., 2019), long-lasting clinical response rates in patients with advanced disease are limited by the development of resistance, of which the mechanisms have not been fully elucidated. Using *K14cre;Brca1^F/F^;Trp53^F/F^*(KB1P) and *K14cre;Brca2^F/F^;Trp53^F/F^* (KB2P) genetically engineered mouse models of hereditary breast cancer (Jonkers et al., 2001; Liu et al., 2007) we have identified various BRCA1- or BRCA2-independent mechanisms of PARPi resistance (reviewed in Gogola et al., 2019). In stark contrast to KB1P tumors (Francica et al., 2020; Gogola et al., 2019), in none of the PARPi-resistant KB2P BRCA2-deficient tumors we found evidence for homologous recombination (HR) restoration (Gogola et al., 2018). Instead, 1/3 of PARPi-resistant KB2P tumors showed a homozygous loss of the poly(ADP-ribose) glycohydrolase (*Parg*) gene. Another resistance mechanism identified in the KB2P model is the protection of replication fork (RF) stability (Ray Chaudhuri et al., 2016), but the molecular mechanism that underlies this alteration in the KB2P model is not yet known.

Regarding the effect of PARPi on RF biology, it has been previously reported that RFs, which are reversed upon DNA damage, are maintained in the reversed state by transient PARP-mediated inhibitory ADP ribosylation of the RECQ1 helicase (Berti et al., 2013; Zellweger et al., 2015). PARP1 therefore acts as a molecular switch to control transient fork reversal and RF restart following genotoxic stress (Zellweger et al., 2015). Whereas untreated cells gain extra time to repair DNA damage through RF reversal, PARPi-treated cells are unable to efficiently maintain forks in a reversed state, resulting in increased DNA breakage and the requirement for HR-mediated DSB repair. Furthermore, the group of Jiri Bartek recently reported that PARPi increases the speed of RF elongation by interfering with the PARP1-p53-p21-fork speed-regulatory network (Maya-Mendoza et al., 2018). This suggests that alterations of RF biology affect the success of PARP inhibition.

Here, we show MDC1 to be an important mediator of the cellular response to PARPi and cisplatin, an effect we link to its participation in the restart of reversed RFs. PARPi-induced uncontrolled fork restart and increased speed of RF progression are counteracted by MDC1 loss, leading to PARPi resistance in BRCA2;p53-deficient mammary tumors. In line with this model, MDC1 loss also causes PARPi resistance in BRCA1-deficient tumor cells without restoring RAD51 foci formation, a typical hallmark of HR restoration that we consistently found in PARPi-resistant KB1P tumors that lose a functional 53BP1-RIF1-shieldin pathway (Barazas et al., 2018; Dev et al., 2018; Jaspers et al., 2013; Noordermeer et al., 2018). Hence, our work reveals a previously unknown function of MDC1 in RF biology, a role that contributes to - DNA damage induced by genotoxic agents and the response of BRCA1- or BRCA2-deficient tumors to clinically relevant anti-cancer drugs, such as PARPi and cisplatin.

## RESULTS

### *Mdc1* depletion causes PARPi resistance *in vitro* and *in vivo*

To identify mechanisms of PARPi resistance in KB2P tumors, we carried out functional genetic screens in the KB2P3.4 cell line, which is derived from a treatment-naïve *Brca2^-/-^;Trp53^-/-^*mouse mammary tumor (Evers et al., 2008). In these cells we introduced pool B of the mouse GeCKO v2 library, a CRISPR/Cas9-based single plasmid system (lentiCRISPRv2) that targets 20,628 mouse genes (Sanjana et al., 2014). The cells were then selected for 3 weeks with 200 nM of the PARPi AZD2461 (Figure 1A), a concentration that kills at least 90% of the parental cells (data not shown). The advantage of AZD2461 is that it is a poor substrate for P-glycoprotein (P-gp), hence the selection of cells with increased P-gp expression is not complicating this assay (O’Connor et al., 2016). Sequencing of the PARPi-surviving populations of 6 biological replicates of this screen revealed a reproducible enrichment of sgRNAs targeting *Mdc1*. The strong effect of *Mdc1* depletion is shown by its high RRA score of 5.78×10^-7^ among all positively selected genes, as determined by the MAGeCK algorithm (Li et al., 2014) (Figure 1A; Table S1).

**Figure 1.**
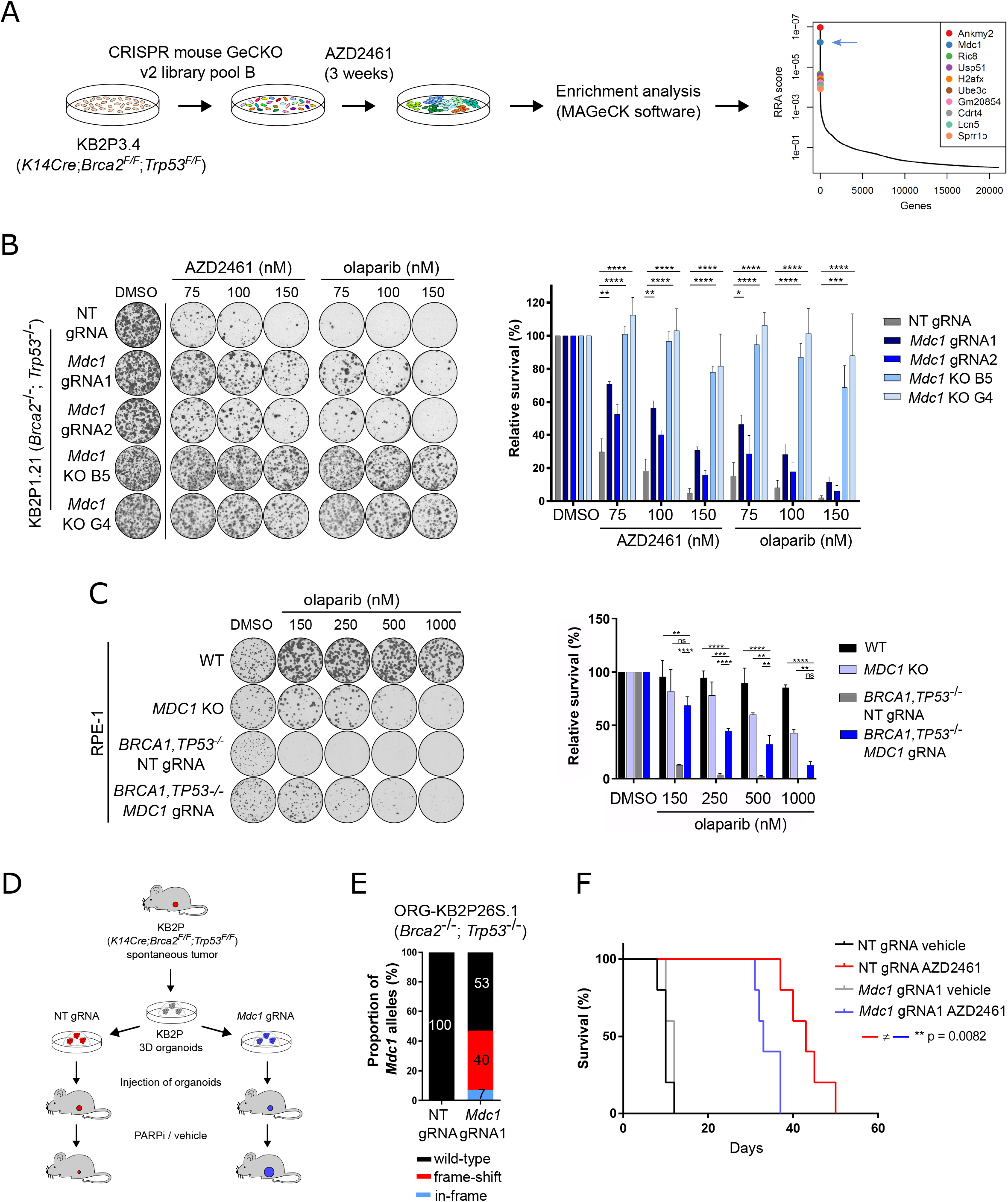
Loss of MDC1 promotes PARPi resistance *in vitro* and *in vivo.* **A)** A schematic overview of the genome-wide loss of function genetic screens. gRNAs are ranked based on *P*-values and robust rank aggregation (RRA) algorithm. **B)** Representative images and quantification of clonogenic assays with KB2P1.21 cells treated with PARP inhibitors or DMSO. The data represent mean ± SEM of three independent experiments. **C)** Representative images and quantification of clonogenic assays with RPE-1 cells treated with olaparib or DMSO. The data represent mean ± SEM of two independent experiments. **D)** *In vivo* validation using genetically modified 3D organoid cultures. **E)** TIDE analysis of 3D organoid lines used in the in vivo experiment. **F)** Kaplan-Meier curve showing the overall survival of the vehicle- or AZD2461-treated mice.

To validate the role of MDC1 in PARPi resistance in BRCA2-defective cells, we used an independent *Brca2^-/-^;Trp53^-/-^* mouse mammary tumor cell line (KB2P1.21) to generate polyclonal KB2P1.21 cell lines by targeting the *Mdc1* gene with two different gRNAs (guide RNA). We then determined the targeting efficacy using the TIDE (Tracking of Indels by Decomposition) analysis (Figure S1A) (Brinkman et al., 2014) and immunofluorescence staining of MDC1 foci formation upon irradiation (Figure S1B and S1C). Both gRNAs resulted in more than 50% of frame-shift modifications and loss of MDC1 foci formation, and we successfully cloned two knockout (KO) cell lines (B5 and G4) with complete absence of MDC1 foci formation (Figures S1A; S1B and S1C). The MDC1-deficient lines showed a significantly improved survival during PARPi exposure, with the clonal lines displaying the strongest resistance phenotype (Figures 1B). The results were corroborated by a significant increase in the frequency of alleles with *Mdc1* frameshift modifications following PARPi treatment in the polyclonal cells (Figure S1D). Of note, the observed PARPi resistance is independent of alterations in PARP1 expression (Figure S1E). We also confirmed the PARPi resistance phenotype in MDC1-deficient derivatives of the KB2P3.4 cell line used for the library screen (Figures S1F; S1G and S1H). Interestingly, MDC1 deficiency also causes PARPi resistance in the BRCA1;p53-deficient background, as demonstrated using the human retinal pigment epithelium RPE-1 cells and mouse mammary tumor-derived KB1P-G3 cells (Figures 1C; S1I; S2A and S2B). Importantly, MDC1 deficiency does not confer resistance to only PARPi, as we also observed a significantly reduced sensitivity to cisplatin (Figure S2E). To investigate whether the PARPi resistance can be recapitulated *in vivo*, we expressed the non-targeting (NT) or *Mdc1*-targeting gRNAs in the KB2P tumor-derived organoids and grafted them orthotopically into a mammary gland of mice as described previously (Duarte et al., 2018) (Figure 1D). Even though the *Mdc1*-targeting efficiency in the organoids was incomplete (40% and 43%, Figures 1E and S2D), we observed a significantly reduced survival of animals bearing the *Mdc1*-targeted KB2P organoid-derived tumors in response to both AZD2461 and olaparib (Figures 1F and S2E). Together, these findings show that loss of MDC1 counteracts the efficacy of PARPi in BRCA1/2-deficient tumor cells both *in vitro* and *in vivo*.

### MDC1 deficiency reduces replication fork velocity and improves the DNA damage tolerance of BRCA2-deficient cells

We and others have previously shown that restoration of HR is a frequent mechanism of chemoresistance (Barazas et al., 2018; Dev et al., 2018; Francica et al., 2020; Jaspers et al., 2013; Noordermeer et al., 2018). Importantly, MDC1;BRCA1;p53-deficient RPE-1 cells did not regain the ability to form RAD51 ionizing radiation-induced foci (IRIF) (Figure 2A). Regarding BRCA2-deficient cells, PARPi resistance has been previously associated with restoration of RF stability (Ray Chaudhuri et al., 2016). Interestingly, MDC1-deficient KB2P1.21 cells did not show this phenotype following HU-induced replication stress (Figure S3A). Instead, we observed a marked decrease in RF progression, even in unperturbed conditions (Figure 2A). While two different polyclonal KB2P1.21 lines expressing independent *Mdc1*-targeting gRNAs displayed only a relatively mild (15-20%) reduction in speed, two independent *Mdc1* knockout clones (B5 and G4) showed a speed reduction of approximately 40% (Figures S3B). We also observed a significant level of RF speed reduction in BRCA1-deficient KB1P-G3 cells expressing *Mdc1*-targeting gRNA (Figure S3C).

**Figure 2.**
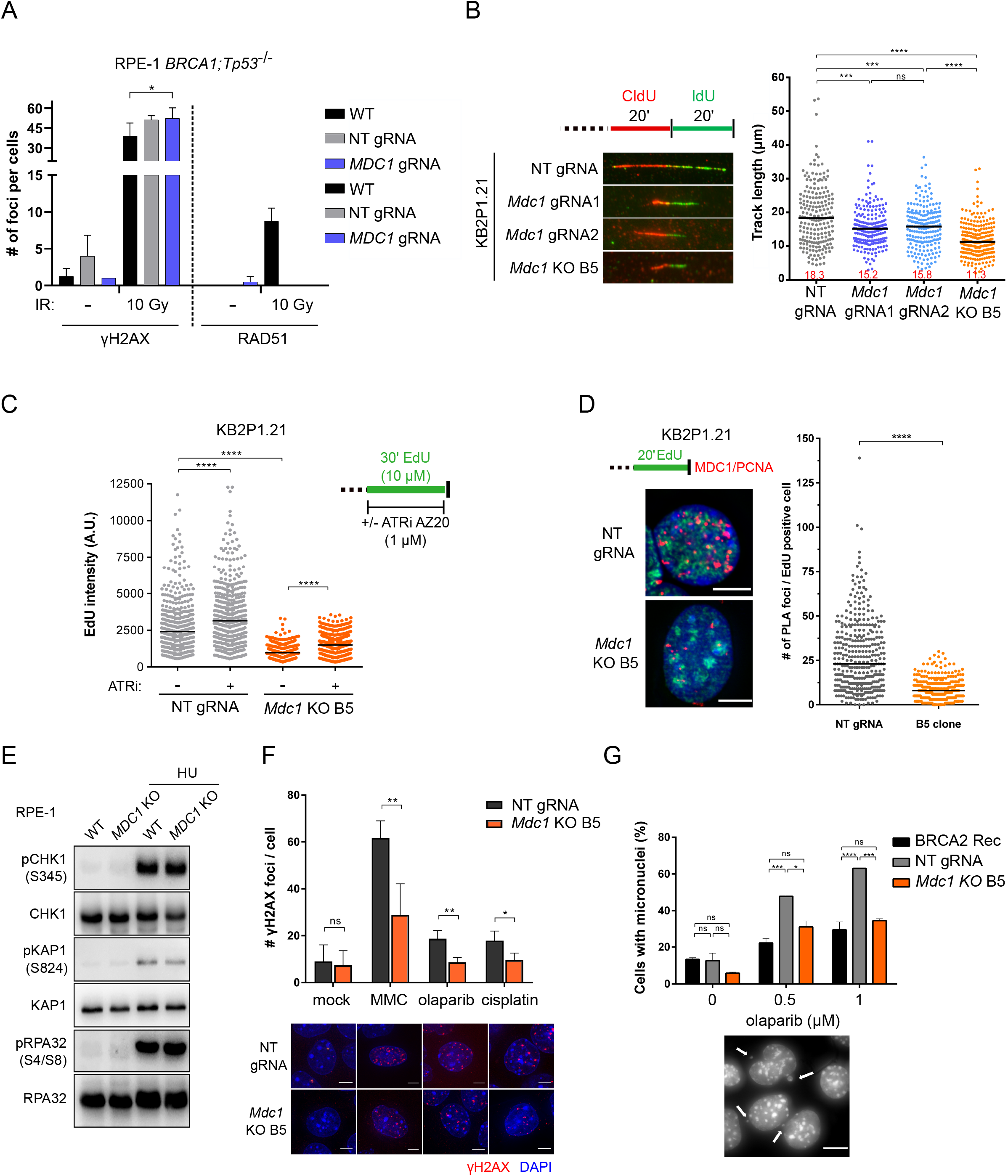
MDC1 deficiency reduces replication fork speed and improves the DNA damage tolerance of BRC2-deficient cells. **A)** Quantification of RAD51 foci formation in RPE-1 cells 4h upon irradiation with 10 Gy. Mean + SD of two independent experiments is shown. **B)** DNA fiber assay in KB2P1.21 cells with representative images of individual fibers. Median of track lengths is shown. Similar results were obtained from 2 independent experiments. **C)** The total replication speed assay. Median of EdU intensity measured in 500 KB2P1.21 cells using a cytometer is shown. Similar results were obtained from 2 independent experiments. **D)** Representative images and analysis of proximity ligation assay showing the interaction of MDC1 with PCNA (red) in EdU-positive cells (green). Median of values from three independent experiments is shown. Scale bars represent 5 µm. **E)** Western blotting of DDR markers before and after 1 mM HU for 24 h in WT and *MDC1* knockout RPE-1 cells. Similar results were observed in two independent experiments. **F)** Quantification and representative images of γH2AX foci in the KB2P1.21 cells after 24 h treatment with DMSO, 300 nM mitomycin C (MMC), 1 µM olaparib or 100 nM cisplatin. Mean ± SD of four independent experiments is shown. The scale bar represents 10 µm. **G)** Percentage of micronuclei positive KB2P1.21 cells after 48 h treatment with DMSO or olaparib. Mean ± SD of values from two independent experiments is shown. A representative image with examples of micronuclei; the scale bar represents 10 µm.

A similar reduction in RF velocity has also been reported in normally growing, BRCA1/2- proficient cells that lost DNA damage response proteins, including RNF8, RNF168 and 53BP1 (Schmid et al. 2018). When we tested the effects of MDC1 loss in BRCA1/2-proficient human RPE-1 cells, we also observed a significant reduction in RF progression, which was fully rescued by WT MDC1 complementation (Salguero et al., 2019) (Figure S3D and S3E). This strongly suggests that the effect of MDC1 on RF speed is BRCA1/2-independent.

MDC1, together with its interacting partner TopBP1, has previously been shown to play a role in activation of the intra-S phase checkpoint upon induction of replication stress (Wang et al., 2011; Wu et al., 2008). To investigate whether the reduced RF velocity observed upon loss of MDC1 is a consequence of unscheduled activation of latent origins of replication due to impaired replication checkpoint control, we assessed the total rate of EdU incorporation in intact nuclei from KB2P1.21 cells (Figures 2C and S4A). The changes in the rates of synthesis at individual RFs and the total DNA synthesis have been previously shown to be inversely correlated upon release of new origins (Petermann et al., 2010; Zhong et al., 2013). Indeed, a significant increase in the EdU intensity was observed in both wild-type (WT) and *Mdc1* KO cells by ATR inhibition, which is known to promote new origin firing (Moiseeva et al., 2017; Shechter et al., 2004). Compared to WT cells, the *Mdc1* KO cells also showed a dramatic reduction in the total EdU intensity in both untreated and ATRi-treated conditions (Figures 2C). These results demonstrate that the observed reduction in RF speed is independent of the latent origin firing. Importantly, the slower replication rate in the *Mdc1* KO KB2P1.21 cells does not correlate with a significant change in the cell proliferation rate (Figure S4B).

Since our results suggest that the effect of MDC1 loss on RF dynamics is independent of a global checkpoint signaling, we performed proximity ligation assays (PLA) and *in situ* analysis of protein interactions at DNA replication forks (SIRF) to determine whether MDC1 physically localizes to active RFs during unperturbed S phase. Indeed, the substantial number of PLA and SIRF foci in MDC1-proficient compared to MDC1-deficient KB2P1.21 and RPE-1 cells demonstrates that MDC1 localizes to active RFs during normal replication (Figure 2D and S4C). Furthermore, the localization of MDC1 at RFs increases following replication stress induced by 2 mM HU (Figure S4C).

To investigate the impact of fork slowing in MDC1-deficient cells on global DNA damage response activation, we investigated typical hallmarks of DNA damage and replication stress, such as phosphorylation of RPA, KAP1 and CHK1 in WT and *MDC1* KO RPE-1 cells (Figure 2E). Consistent with the depletion of another DSB repair factor, RNF168 (Schmid et al., 2018), slower RF progression during normal S phase in MDC1-depleted cells is not associated with replication stress and the global DDR activation (Figure 2E). Similar results were obtained in *Mdc1* KO BRCA2;p53-deficient KB2P1.21 cells (Figure S4D). Interestingly, 24h treatment of MDC1-deficient KB2P1.21 cells with the replication-targeting agents mitomycin C (MMC), olaparib or cisplatin caused an approximately 50% reduction in the number of the γH2AX foci, relative to MDC1-proficient KB2P1.21 cells (Figure 2F). This cannot be explained by a reduced ability of cells to form γH2AX foci in the absence of MDC1 (Figures 2A and S3E). Compared to MDC1-proficient KB2P1.21 cells, MDC1-deficient cell cultures also showed significantly fewer cells with micronuclei following 48h exposure to olaparib (Figure 2G). These data show that slower RF progression in BRCA2-deficient cells with MDC1 loss does not impose a threat to genome integrity in unperturbed conditions. Under stress conditions, the loss of MDC1 even improves the tolerance to DNA damage.

### Loss of MDC1 delays replication fork restart and leads to an accumulation of reversed forks

Since the reduction in RF fork speed does not occur as a consequence of an unscheduled release of latent origins, we investigated the role of MDC1 in the dynamics of fork remodeling. It has been previously shown that RF reversal is a highly dynamic process enabling cells to attenuate the cytotoxic effects of a wide range of replication stress-inducing stimuli of endogenous and exogenous origin (Neelsen and Lopes, 2015; Zellweger et al., 2015). Dysregulation of the reversed fork metabolism or their restart into the classical three-way junctions have been shown to result in an altered fork progression rate (Bennett et al., 2020; Berti et al., 2013; Rainey et al., 2020; Raso et al., 2020). To investigate whether the reduced RF speed in MDC1-deficient cells is accompanied by an increased frequency of reversed RF intermediates, we quantified RFs during normal replication using electron microscopy (Figure 3A). Indeed, while we detected only 5% of the reversed fork intermediates in the WT RPE-1 cells, an approximately 3-fold higher frequency was observed in the *MDC1* KO cells (Figure 3A). A similar level of reversed fork frequency upon loss of other DDR factors has previously been reported (Schmid et al., 2018). To test whether the increased reversed fork frequency can be explained by altered remodeling dynamics, we assessed thecapacity of KB2P1.21 cells to restart forks following a mild 2 mM hydroxyurea (HU) treatment (Figures 3B and 3C). Indeed, a significantly lower frequency of restarted forks was detected in the MDC1-deficient KB2P1.21 clones B5 and G4 40 min after HU removal relative to MDC1-proficient cells. Interestingly, this difference disappeared after 80 min of recovery following the HU release, which shows that the delay in fork restart in MDC1-deficient cells is only temporary (Figures 3B and 3C). These results indicate that MDC1 is important for the regulation of RF restart upon replication stress-induced fork reversal.

**Figure 3.**
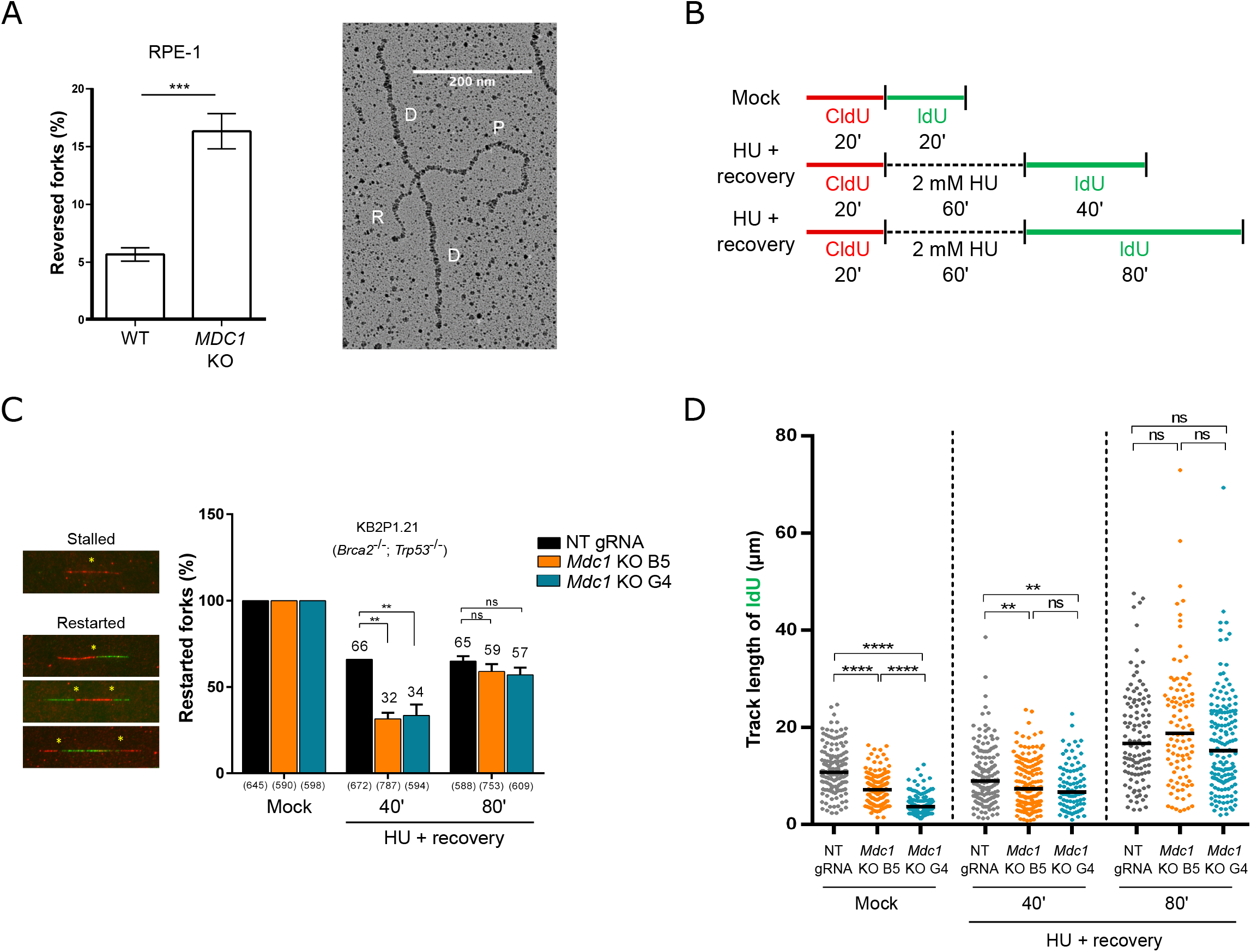
Loss of MDC1 delays replication fork restart and leads to an accumulation of reversed forks. **A)** Frequency of reversed RFs in untreated WT or *MDC1* knockout RPE-1 cells analyzed by transmission electron microscopy. Mean ± SD of three independent experiments is shown. An electron micrograph of a representative reversed fork. **B)** Scheme of RF restart experiments. **C)** Examples of stalled or restarted forks, the asterisks show examples of the quantified events. Analysis of the percentage of restarted forks without treatment or after RF stalling with HU and two recovery time-points. Mean ± SD of two independent experiments is shown. The numbers of counted RFs are shown in brackets. **D)** IdU track length analysis of the restarted forks. Median of values from two independent experiments is shown.

To gain more insight into the underlying mechanism of the delayed recovery of stalled RFs, we measured the track length of the green IdU label given after mock or HU treatment respectively (Figure 3D). As expected, significantly shorter IdU tracks were measured in the native condition in both MDC1-deficient KB2P1.21 lines B5 and G4. Surprisingly, we measured a much smaller difference between the MDC1-proficient and MDC1-deficient lines 40 min after HU release (Figure 3D). Together with the observed decrease in the frequency of restarted forks at the 40 min time-point, this finding suggests the existence of an alternative, MDC1-independent fork restart pathway in BRCA2-deficient cells. The lack of a statistically significant difference in the IdU track lengths at the 80 min time-point further demonstrates that cells are not fully dependent on MDC1 to efficiently restart stalled replication forks (Figure 3D). Furthermore, the data suggest that while the restart of RFs following HU-mediated stalling is delayed in the MDC1,BRCA2-deficient cells, their progression rate may increase relatively to the control BRCA2-deficient cells (Lemaçon et al., 2017). This difference may not be detectable in the early 40 min time-point due to the overall lower rate of DNA synthesis (Figure 3D).

Together, these data demonstrate a new role of MDC1 in the regulation of restart of reversed RFs under both unperturbed and replication stress conditions.

### MDC1 deficiency reduces replication fork speed and promotes PARPi resistance by regulating MRE11 activity at the reversed forks

PARP1 has been shown to stabilize RFs in the regressed state and regulate their restart by limiting the activity of the RECQ1 helicase (Berti et al., 2013). Using a high PARPi concentration of 10 μM olaparib – a 100-fold higher concentration than used in our clonogenic survival assays (Figures 1B; 1C; S1F and S2B) – this inhibitory effect of PARP1 on RECQ1-mediated RF restart can be bypassed (Berti et al., 2013). Indeed, treatment with 10 μM olaparib restored the fork speed in MDC1-deficient mouse KB2P1.21 and human RPE-1 cells (Figures 4A and 4B). This further supports our notion that reduced RF progression in MDC1-deficient cells is caused by delayed fork restart. Of note, the effect of PARPi on RECQ1-mediated fork restart is dose-dependent and does not occur following treatment with 1 μM olaparib (Figure 4C), a concentration that is still substantially higher than what is needed to kill KB2P cells (Figures 1B and S1F). Consistent with this observation, no effect on replication speed was observed after a 1-week long olaparib exposure with the dose used in our *in vitro* clonogenic assays (Figure 4D).

**Figure 4.**
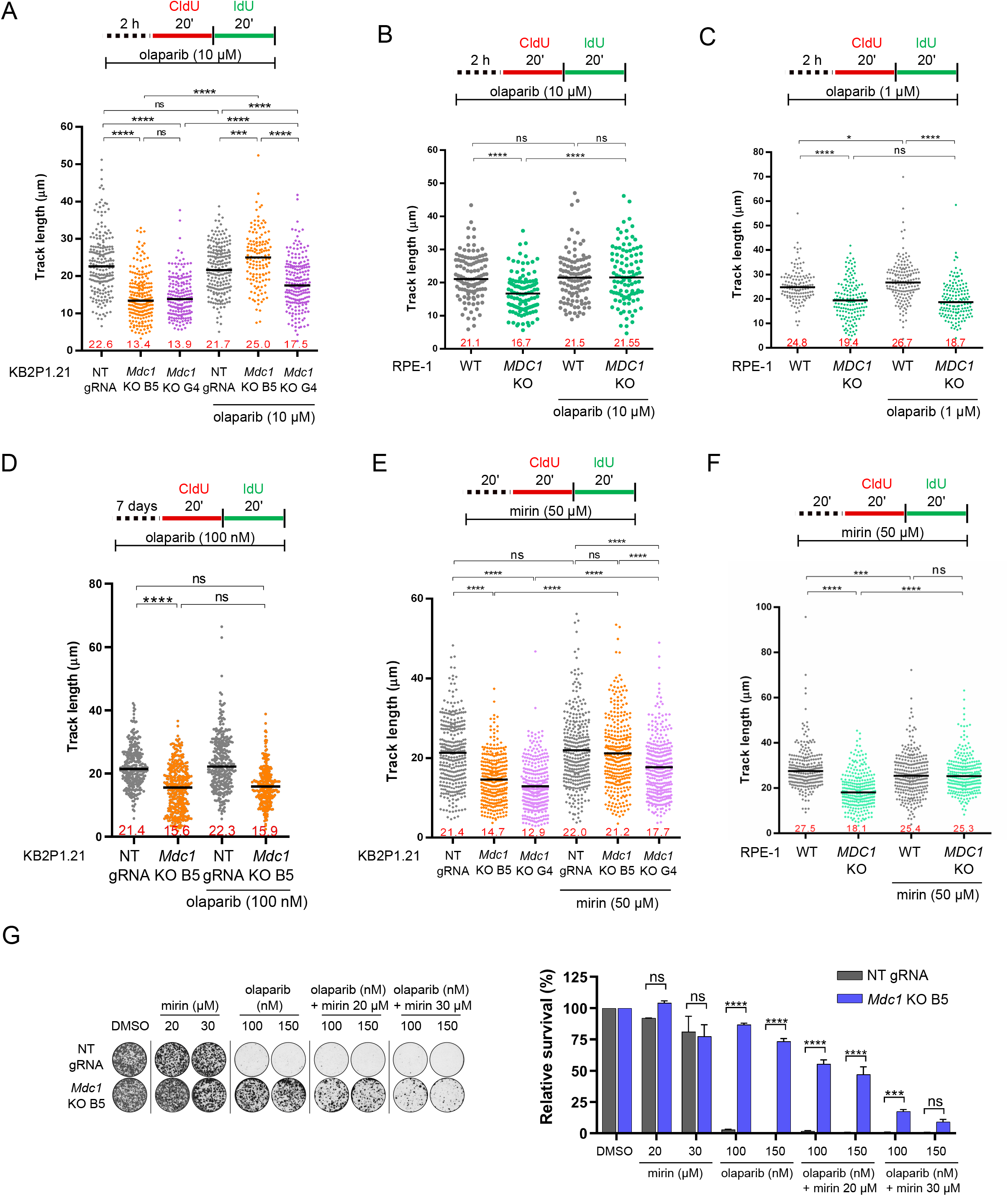
MDC1 regulates replication fork speed and PARPi sensitivity by suppressing MRE11 activity at the reversed forks. **A)** Track length analysis in KB2P1.21 cells after treatment with DMSO or 10 µM olaparib. Median of two independent experiments is shown. **B-C)** Track length analysis in RPE-1 cells after treatment with DMSO, 1 µM or 10 µM olaparib. Median of at least 100 fork lengths is shown. Similar results were obtained in two independent experiments. **D)** Track length analysis in KB2P1.21 cells after 7-day treatment with 100 nM olaparib. Median of two independent experiments is shown. **E-F)** Total track length in KB2P1.21 (D) or RPE-1 (E) cells without and after treatment with 50 µM mirin. Median of two independent experiments is shown. **G)** Representative images (left) and quantification (right) of clonogenic assay with KB2P1.21 cells showing treatment response to mirin or olaparib alone, and to a combined treatment. Mean ± SD of two independent experiments is shown.

In previous studies, it was shown that MRE11-mediated processing of reversed RFs during unperturbed S phase affects the rate of fork progression (Bennett et al., 2020; Schmid et al., 2018). We therefore hypothesized that MDC1 antagonizes this function of MRE11. To test whether MRE11-mediated processing is implicated in the reduced RF dynamics in MDC1-deficient cells, we performed the DNA fiber assay in the KB2P1.21 and RPE-1 cell lines in the presence of the MRE11 exonuclease activity inhibitor mirin. Indeed, MRE11 inhibition resulted in the full rescue of the fork progression rate in the MDC1-deficient KB2P1.21 and RPE-1 cells (Figures 4E and 4F). MRE11 inhibition may actually serve as an approach to re-sensitize MDC1-depleted tumors to PARPi. While continuous treatment with mirin alone did not have a clear effect on cell survival, we observed a dose-dependent sensitization of MDC1-deficient cells to olaparib when combined with mirin (Figure 4G). Hence, PARPi resistance caused by reduced replication speed and subsequent DNA damage tolerance of MDC1-depleted cells can be counteracted by blocking MRE11.

## Discussion

In this study, we identified loss of the DSB repair protein MDC1 as a novel mechanism causing PARPi resistance in BRCA1/2-deficient cells *in vitro* and in BRCA2-deficient tumors *in vivo*. In addition to their resistance to PARPi, MDC1-deficient cells were also more tolerant to damage induced by other anti-cancer drugs, including cisplatin and MMC.

While the role of MDC1 in canonical DSB repair is well-known, its function in RF metabolism was unknown. Using iPOND assays following replication stress, the group of David Cortez found MDC1 to be present at RFs (Sirbu et al., 2013). Here, we show that MDC1 is already present at RFs of normally growing cells, where it contributes to RF restart. Mechanistically, MDC1 appears to counteract MRE11-mediated processing of reversed RFs. While other DDR factors, such as RNF8, RNF168 and 53BP1 have previously been shown to modulate RF progression by limiting nucleolytic processing of the reversed forks (Schmid et al., 2018), it is not known how loss of these factors affects chemotherapy response in the context of the BRCA1/2 deficiency. Thus far, genetic alterations resulting in defective RF progression, stability or restart have usually been linked to an elevated genome instability and increased sensitivity of cells to replication stress or DNA damage-inducing agents (Bennett et al., 2020; Her et al., 2018; Mukherjee et al., 2019; Rainey et al., 2020; Schmid et al., 2018; Xu et al., 2017). In this study, we show that loss of MDC1 can also have the opposite effect. The delayed fork restart and reduced fork speed in MDC1-deficient cells does not stimulate the activation of global DDR signaling markers. In contrast, it results in an improved DNA damage tolerance and chromosomal stability in the context of BRCA1/2 deficiency. Our findings are consistent with the recent observation of an ATR-mediated global fork reversal as a cellular strategy to limit potentially deleterious collisions of active RFs with DNA lesions, such as inter-strand cross-links (Mutreja et al., 2018).

Furthermore, we show that rescue of the normal fork dynamics with the MRE11 inhibitor mirin re-sensitized MDC1-deficient KB2P1.21 cells to the PARPi olaparib. While extensive MRE11-mediated degradation of reversed forks in BRCA1/2-defective cells lacking proper fork stabilization capacity has been previously associated with their increased sensitivity to chemotherapeutic agents (Ray Chaudhuri et al., 2016; Mijic et al., 2017), loss of MDC1 seems to result in an MRE11-induced lower fork speed. Importantly, while we observed HU-mediated RF degradation in MDC1-deficient cells, the cell survival data do not support increased RF collapse and chromosomal instability following treatments with PARPi or cisplatin.. The amount of MRE11-mediated resection and its spatio-temporal dynamics may therefore be the critical determinant of PARPi and cisplatin response of BRCA-deficient tumors. Thus, the complex nature of the nucleolytic processing at unchallenged or stalled RFs requires more research to be fully understood.

In summary, we show that MDC1 loss represents a novel mechanism of chemoresistance in BRCA1/2-deficient cells, which is independent of rewiring homology-directed DNA repair or restoration of RF protection. Instead, remodeling of the MDC1-mediated RF restart drives PARPi resistance via increased DNA damage tolerance.

## Acknowledgments

We wish to thank Piet Borst, Carmen Disler and Horst Posthaus for critical reading of the manuscript. Moreover, we thank the members of the Preclinical Intervention Unit of the Mouse Clinic for Cancer and Ageing (MCCA) at the Netherlands Cancer Institute (NKI), Georgina Lackner and Fabiana Steck for their help at the Vetsuisse Faculty mouse facility and Denise Howald for her technical support. Financial support came from the Swiss National Science Foundation (310030_179360 to S.R.), the European Research Council (CoG-681572 to S.R. and H2020-MSCA-IF-2016-743290 to J.S.B.), the Swiss Cancer League (KLS-4282-08-2017 to S.R.), the Boehringer Ingelheim Fonds PhD fellowship (to M.L.), and the Wilhelm-Sander Foundation (no. 2019.069.1 to S.R.).

## Author Contributions

Conceptualization, M.L., J.S.B., E.G., J.J. and S.R.; Methodology, M.L., J.S.B., E.G., A.A.D., D.D., J.S., I.S., M.v.d.V., S.P.J., M.L., J.J., and S.R.; Investigation, M.L., J.S.B., E.G., A.A.D., D.D., J.S., I.K., M.M., M.S., P.F., and R.d.K.-G.; Software and Formal Analysis, M.L., J.S.B., and E.G.; Resources, M.S., M.v.d.V., S.P.J., M.L., J.J. and S.R.; Writing, M.L. and S.R.; Supervision, M.v.d.V., M.L., J.J., and S.R.; Funding Acquisition, M.L., J.S.B. and S.R.

## Declaration of Interests

The authors declare no competing interests.

## STAR METHODS

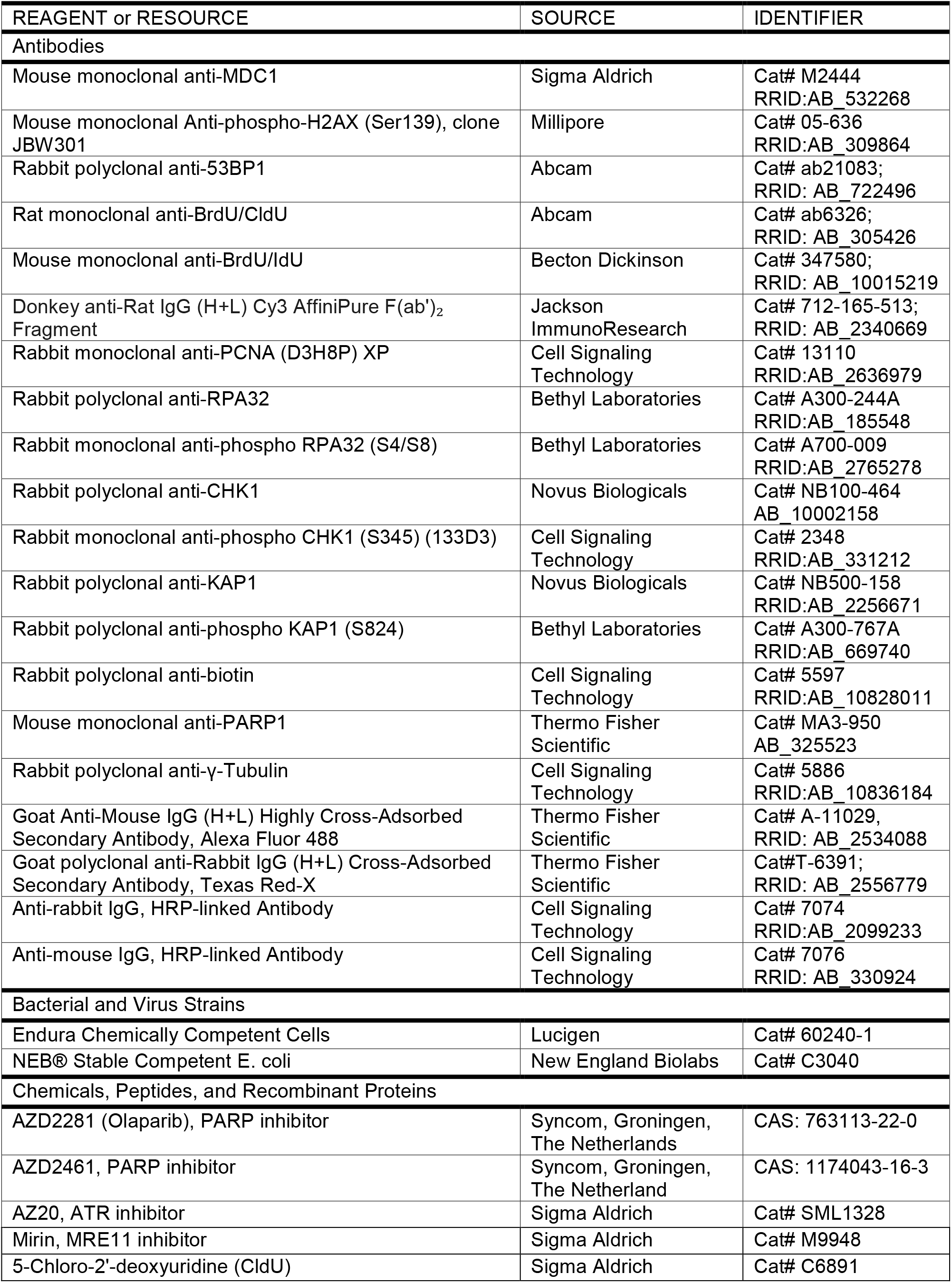
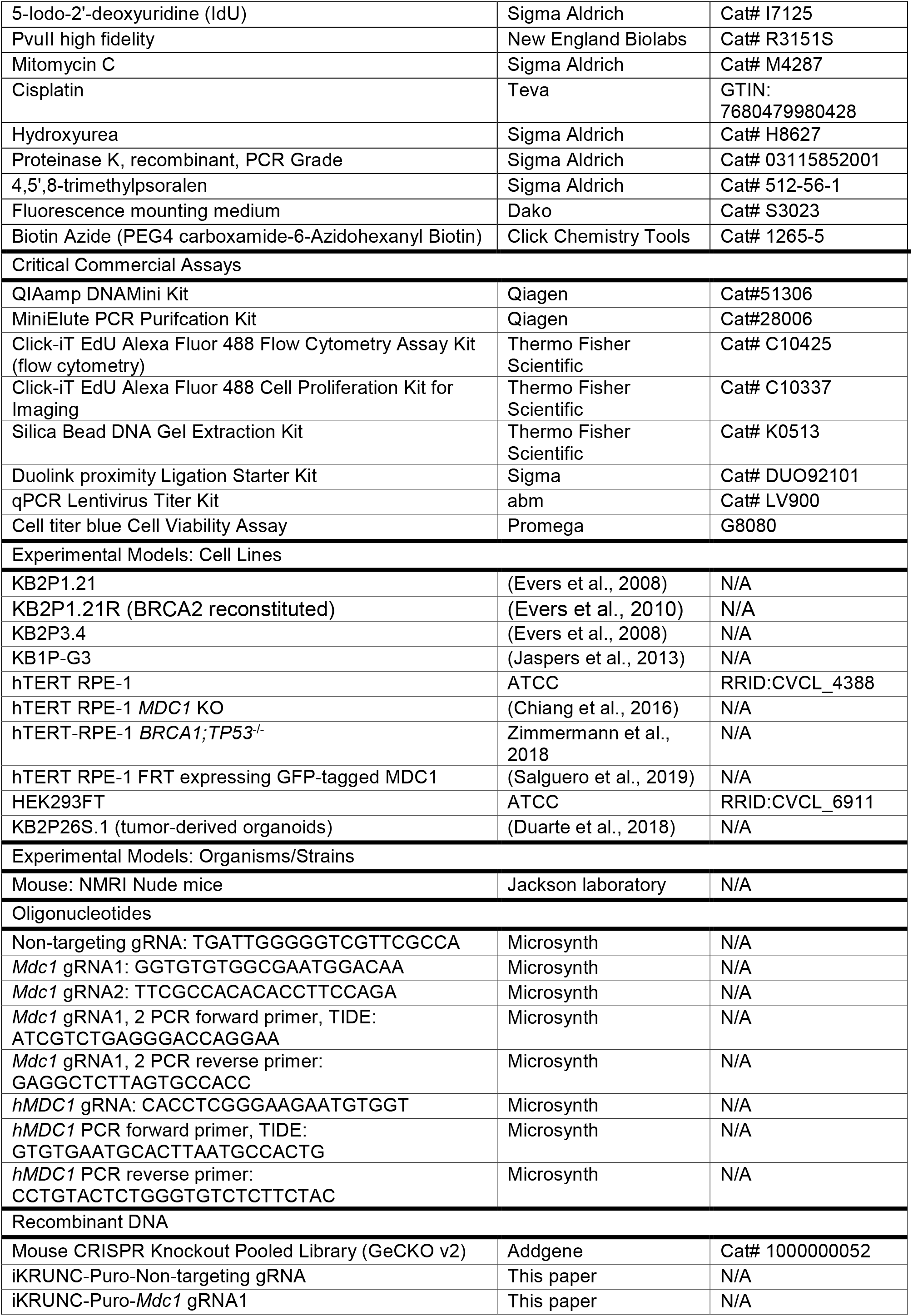
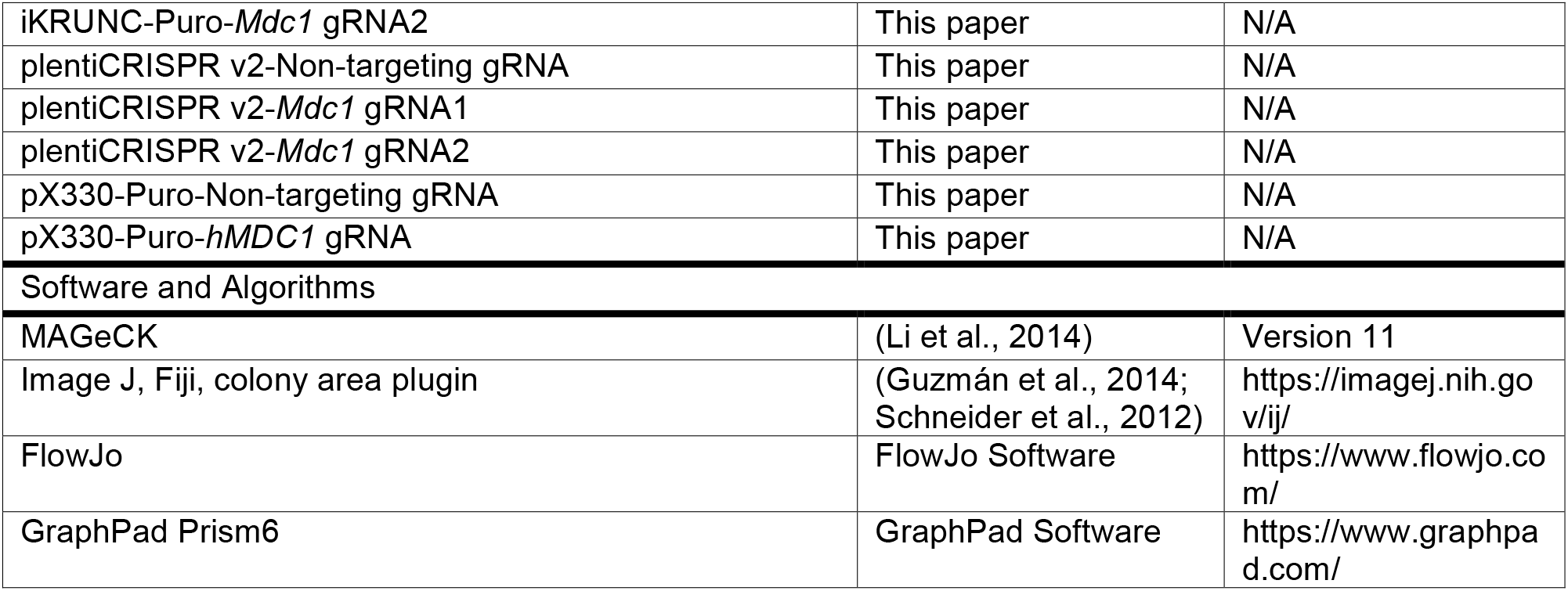
KEY RESOURCES TABLE

## RESOURCE AVAILABILITY

### LEAD CONTACT

Further information and requests for resources and reagents should be directed to and will be fulfilled by the Lead Contact, Sven Rottenberg.

### MATERIALS AVAILABILITY

All unique/stable reagents generated in this study will be made available on request, but we may require a payment and/or a completed Materials Transfer Agreement if there is potential for commercial application.

### DATA AND CODE AVAILABILITY

Sequencing of the CRISPR/Cas9 genetic screens was performed at the Netherlands Cancer Institute. The results of the MAGeCK analysis are available in Table S1.

### LEAD CONTACT AND MATERIALS AVAILABILITY

Further information and requests for resources and reagents should be directed to and will be fulfilled by the Lead Contact, Sven Rottenberg.

### EXPERIMENTAL MODEL AND SUBJECT DETAILS

#### Mice

The Animal Ethics Committee of The Netherlands Cancer Institute (Amsterdam, the Netherlands) and the Animal Ethics Committee of the Canton of Bern (Switzerland, Application number BE40/18) approved all animal experiments. All experiments were performed in accordance with the Dutch Act on Animal Experimentation (November 2014) and the Swiss Act on Animal Experimentation (December 2015). CRISPR-Cas9-modified organoids lines derived from *K14cre;Brca2^F/F^;Trp53^F/F^* (KB1P) female mice were transplanted in 6-9 weeks-old NMRI nude mice for the *in vivo* validation.

#### Cell Lines

The KB2P1.21 and KB2P3.4 cell lines were previously established from a *K14cre;Brca2^F/F^;Trp53^F/F^* (KB2P) mouse mammary tumor as described by Evers et al., 2008. The KB2P1.21R cell line was derived from the KB2P1.21 cells by reintroducing BRCA2 as described by Evers et al., 2010. The KB1P-G3 cell line was previously established from a KB1P mouse mammary tumor and cultured as described by Jaspers et al., 2013. The hTERT-RPE-1 *MDC1* knockout cell line and hTERT-RPE-1 FRT-derived cell line stably expressing inducible WT MDC1 GFP-tagged construct were established and previously described by Salguero et al., 2019. The hTERT-RPE-1 *BRCA1;TP53*^-/-^ cells were a kind gift from Daniel Durocher. The cell line was established and previously described by Zimmermann et al., 2018. All the KB1P and KB2P lines were grown in Dulbecco’s Modified Eagle Medium/Nutrient Mixture F-12 (DMEM/F12; Gibco) supplemented with 10% fetal calf serum (FCS, Sigma Aldrich), 50 units/ml penicillin-streptomycin (Gibco), 5 µg/ml Insulin (#I0516, Sigma Aldrich), 5 ng/ml cholera toxin (#C8052, Sigma Aldrich) and 5 ng/ml murine epidermal growth-factor (EGF, #E4127, Sigma Aldrich). The HEK293FT cell line (RRID:CVCL_6911) was cultured in Iscove’s Modified Dulbecco’s Media (IMDM, Gibco) supplemented with 10% fetal calf serum (FCS, Sigma Aldrich) and 50 units/ml penicillin-streptomycin (Gibco). All the hTERT-RPE-1 cell lines were cultured in Dulbecco’s Modified Eagle Medium/Nutrient Mixture F-12 (DMEM/F12; Gibco) supplemented with 10% FCS, 100 U/ml penicillin, 100 ng/ml streptomycin, 17 ml NaHCO3 7.5% per 500 ml (Sigma Aldrich) and 2 mM L-glutamine. In addition, medium of the hTERT-RPE-1 FRT derived cells expressing GFP-tagged MDC1 construct contained 0.5 mg/ml G418 disulfate salt solution (#G8168, Sigma Aldrich). Expression of the construct was induced by adding 0.5 µg/ml of doxycycline in the growth medium 48 h prior experiments. The cells were maintained in the presence of doxycycline for the whole duration of the experiment.

Mouse mammary tumor-derived KB2P and KB1P cell lines and the human *BRCA1;TP53*^-/-^ hTERT-RPE-1 cell lines were grown in low oxygen atmosphere at standard temperature (37°C, 3% O2, 5 % CO2). HEK293FT and hTERT-RPE-1 cell lines were grown in standard conditions (37°C, 5% O2 5 % CO2). Testing for mycoplasma contamination was performed on a regular basis.

#### Tumor-Derived Organoids

The KB2P26S.1 3D tumor organoid line was previously established from a *Brca2^-/-^;Trp53^-/-^* mouse mammary tumor and cultured as described by (Duarte et al., 2018). Briefly, cultures were embedded in Culturex Reduced Growth Factor Basement Membrane Extract Type 2 (BME, Trevigen; 40 μl BME:growth media 1:1 drop in a single well of 24-well plate) and grown in Advanced DMEM/F12 (AdDMEM/F12, Gibco) supplemented with 1 M HEPES (Sigma Aldrich), GlutaMAX (Gibco) 50 units/ml penicillin-streptomycin (Gibco), B27 (Gibco), 125 μM N-acetyl-L-cysteine (Sigma Aldrich) and 50 ng/ml murine epidermal growth factor (Sigma Aldrich). Organoids were cultured under standard conditions (37°C, 5% CO2) and regularly tested for mycoplasma contamination.

Further *in vitro* culture details and gene editing details are provided in the method details section.

### METHOD DETAILS

#### CRISPR/Cas9-based genetic screen

The PARPi resistance screens were performed in the KB2P3.4 tumor cell line, which was previously established from a KB2P tumor (Evers et al., 2008). Mouse GECKO v2 library, pool B (62,804 gRNAs targeting 20,628 genes (3 gRNAs/gene) and including 1,000 control non-targeting gRNAs), was stably introduced into the cells by lentiviral transduction at the multiplicity of infection (MOI) of 1.5. 6 independent transductions were carried out to obtain mutagenized cells for biological replicates of the PARPi resistance screen. To perform the genetic screen at 50x library coverage, 3×10^6^ mutagenized KB2P3.4 cells in each replicate were plated in 10-cm flasks, at low density (30,000 cells per flask) and grown in the medium containing 200-300 nM AZD2461 for 3 weeks. The medium with the PARPi was refreshed twice a week. Cells were harvested before and after PARPi treatment for genomic DNA isolation. Subsequently, gRNA sequences were amplified from genomic DNA by two rounds of PCR amplification as described previously (Sanjana et al., 2014; Xu et al., 2015). Resulting PCR products were purified using MinElute PCR Purification Kit (Qiagen) and submitted for Illumina sequencing. Sequence alignment and enrichment analysis (day 0 vs PARPi-treated population) was carried out using MAGeCK software (Li et al., 2014).

#### Gene editing, silencing, plasmids and cloning

##### Lentiviral transductions

Lentiviral stocks were generated by transient transfection of HEK293FT cells. On day 0, 6×10^6^ HEK293FT cells were seeded in 150 cm cell culture dishes and on the next day transiently transfected with lentiviral packaging plasmids and the pGS-Cas9 (Neo) or iKRUNC-Puro / plentiCRSIPRv2 vector containing the respective gRNA or a non-targeting gRNA using 2x HBS (280 nM NaCl, 100 mM HEPES, 1.5 mM Na2HPO4, pH 7.22), 2.5 M CaCl2 and 0.1x TE buffer (10 mM Tris pH 8.0, 1 mM EDTA pH 8.0, diluted 1:10 with dH2O). After 30 h, virus-containing supernatant was concentrated by ultracentrifugation at 20,000 rpm for 2 h in a SW40 rotor and the virus was finally resuspended in 100 μl PBS. The virus titer was determined using a qPCR Lentivirus Titration Kit (#LV900, Applied Biological Materials). For lentiviral transduction, 150,000 target cells were seeded in 6-well plates. 24 h later, virus at the MOI of 50 was applied with 8 μg/ml Polybrene (Merck Millipore). Virus-containing medium was replaced with medium containing puromycin (3.5 μg/ml, Gibco) 24 h later. Puromycin selection was performed for three days; subsequently cells were expanded and frozen down at early passage. Tumor-derived organoids were transduced according to a previously established protocol (Duarte et al., 2018). The target sites modifications of the polyclonal cell pools were analyzed by TIDE analysis which is described below.

##### Genome editing

For CRISPR/Cas9-mediated genome editing using the iKRUNC system described previously by Prahallad et al., 2015, KB2P1.21 and KB2P3.4 cells, or KB2P26S.1 organoids were first transduced with the lentiviral pGS-Cas9 (Neo) construct and grown under G418 selection (500 μg/ml) for 5 days. Next, neomycin-selected cells were incubated with lentiviral supernatants of iKRUNC-Puro vectors containing the respective gRNA or a non-targeting gRNA and exposed to 3 μg/ml puromycin for 5 days. To induce gRNA expression, puromycin-surviving cells were treated for another 5 days with 3 μg/ml doxycycline (Sigma Aldrich). For CRISPR/Cas9-mediated genome editing with lentiCRISPRv2 system, KB1P-G3 cells were transduced with the pLentiCRISPRv2 vector encoding non-targeting gRNA, *Mdc1*-targeting gRNA1 or *Mdc1*-targeting gRNA2. The cells were then grown under Puromycin (3 μg/ml) selection for 5 days. All constructs sequences were verified by Sanger sequencing. gRNA sequences are provided below.

CRISPR gRNA sequences for modification of *Mdc1* were chosen from the GeCKo library v2 (Sanjana et al., 2014). The gRNA sequences were as follows:

*Mdc1* gRNA1: 5’- GGTGTGTGGCGAATGGACAA -3’ targeting exon 4

*Mdc1* gRNA2: 5’- TTCGCCACACACCTTCCAGA -3’ targeting exon 4

Non-targeting (NT) gRNA: 5’-TGATTGGGGGTCGTTCGCCA-3’

For CRISPR/Cas9-mediated genome editing of the *BRCA1;TP53*^-/-^ hTERT-RPE-1 cells, cells were transduced with the pX330 vector encoding non-targeting gRNA or *MDC1*-targeting gRNA. The cells were then grown under Puromycin (3 μg/ml) selection for 5 days. All constructs sequences were verified by Sanger sequencing. gRNA sequences are provided below.

*MDC1* gRNA: 5’- CACCTCGGGAAGAATGTGGT -3’

Non-targeting (NT) gRNA: 5’-TGATTGGGGGTCGTTCGCCA-3’

*gDNA isolation, amplification and TIDE analysis*

To assess the modification rate at the gRNA-targeted region of *Mdc1*/*MDC1*, cells were pelleted and genomic DNA was extracted using the QIAmp DNA mini kit (Qiagen) according to manufacturer’s protocol. Target loci were amplified using Phusion High Fidelity Polymerase (Thermo Scientific) using a 3-step protocol. For amplification of the gRNA target sites in mouse cells:(1) 98°C for 30 s, (2) 35 cycles at 95 °C for 15 s, 55 °C for 15 s and 72 °C for 30 s, (3) 72 °C for 7 min. For amplification of the gRNA target sites in human cells: (1) 98°C for 30 s,

(2) 35 cycles at 95 °C for 15 s, 65 °C for 15 s and 72 °C for 15 s, (3) 72 °C for 7 min. Reaction mix consisted of 10 ul of 2x Phusion Mastermix (Thermo Fisher), 1ul of 10 uM forward and reverse primer and 100 ng of DNA in 20 ul total volume. PCR products were purified using the QIAquick PCR purification kit (Qiagen) according to manufacturer’s protocol and submitted with corresponding forward primers for Sanger sequencing to confirm target modifications using the TIDE algorithm (Brinkman et al., 2014). The primers used in this PCR are listed in Table S2.

#### Clonogenic assays

To assess the growth and survival upon exposure to PARPi, KB2P1.21, KB2P3.4 or KB1P-G3 cells were seeded in 6-well plates in the following densities: 5,000 cells/well (KB2P1.21 and KB2P3.4) and 4,500 cells/well (KB1P-G3). Clonogenic assays with RPE-1 cells were performed by plating 500 cells in 10 cm dish. The treatment of cells with DMSO or indicated concentrations of PARPi olaparib or AZD2461 started at the day of plating the cells and lasted for the whole duration of the experiment. The medium with DMSO or PARPi was refreshed twice a week. In the clonogenic assays with cisplatin, cells were grown in the cisplatin-containing medium for the whole duration of the experiment and the medium was refreshed twice a week. The control, DMSO-treated plates were fixed 7 days (mouse cells) or 10 days (RPE-1 cells) after seeding. The PARPi- and cisplatin-treated plates were fixed after 10 (mouse cells, PARPi) or 14 (mouse cells, cisplatin; RPE-1 cells, PARPi) days. The fixation was done with 4% formalin and the surviving colonies stained with 0.1% crystal violet. The survival and growth of mouse cells was analyzed in an automated manner using the ImageJ ColonyArea plugin described previously (Guzmán et al., 2014). The survival of RPE-1 cells was assessed by manual counting of colonies. For the competition assays, cells were collected before and after the experiment for gDNA isolation and TIDE analysis as described above. For the clonogenic assay with combined olaparib and mirin treatment, 5,000 KB2P1.21 cells were seeded in 6 well plates one day before start of the treatment. Next day, medium containing the indicated concentrations of DMSO, olaparib/DMSO, mirin/DMSO or olaparib/mirin was added to the cells. Medium with drugs was refreshed twice a week. The next steps were carried out as described above.

#### CTB proliferation assay

To analyze the proliferation rate of control (non-targeting gRNA) or *Mdc1*-modified KB2P1.21 cells, 1,000 cells were seeded in 96-well plates one day before the experiment. Rate of proliferation was analyzed using CellTiter-Blue® Cell Viability Assay (Promega) following manufacturer’s instructions. Fluorescence intensity of culture medium upon incubation with CellTiter-Blue reagent was measured on four consecutive days.

#### *In vivo* studies

For tumor organoid transplantation, organoids were collected, incubated with TripLE at 37°C for 5 min, dissociated into single cells, washed in PBS, resuspended in tumor organoid medium and mixed in a 1:1 ratio of tumor organoid suspension and BME. Organoid suspension containing a total of 5×10^4^ cells were injected in the fourth right mammary fat pad of 6-9 week-old NMRI nude mice. Mammary tumor size was measured by caliper on at least three days per week and tumor volume was calculated (length x width^2^ /2). Treatment of tumor bearing mice was initiated when tumors reached a size of ∼75mm^3^, at which point mice were separated into two vehicle-treated groups (NT gRNA n= 5, *Mdc1*-targeting gRNA1 n= 5) and AZD2461-treated groups (NT gRNA n= 5, *Mdc1*-targeting gRNA1 n= 5). AZD2461 (100 mg/kg) was administered orally for 28 consecutive days, control mice were dosed with vehicle only. Animals were anesthetized with isoflurane, sacrificed with CO2 followed by cervical dislocation when the tumor reached a volume of 1,500 mm^3^. Tumor sampling included cryopreserved tumor pieces, fresh frozen tissue and formalin-fixed material (4% (v/v) formaldehyde in PBS). For the *in vivo* experiment using olaparib presented in Figure S2, treatment of tumor bearing mice was initiated when tumors reached a size of ∼150mm^3^, at which point mice were separated into two untreated/vehicle groups (NT gRNA n= 10, *Mdc1*-targeting gRNA1 n= 10) and olaparib-treated group (NT gRNA n= 10, *Mdc1*-targeting gRNA1 n= 10). Treatment with vehicle/olaparib (100 mg/kg) was administered intraperitoneally for 56 consecutive days. Animals were anesthetized with isoflurane, sacrificed with CO2 followed by cervical dislocation when the tumor reached a volume of 1,000 mm^3^.

#### Immunofluorescence

Cells were seeded on coverslips in 24-well plates 3 days before the experiment. For analysis of MDC1 IRIF in KB2P1.21 cells, DNA damage was induced by γ-irradiation (10 Gy) 3 h prior to fixation. Subsequently, cells were washed in PBS and fixed with 4% (v/v) PFA/PBS for 20 min at room temperature (RT). Fixed cells were washed with PBS and were permeabilized for 20 min in 0.2% (v/v) Triton X-100/PBS. Subsequently, slides were washed three times with 0.2% Tween-20/PBS and blocked with staining buffer (PBS, BSA (2% w/v), glycine (0.15% w/v), Triton X-100 (0.1% v/v)) for 1 h at RT. Incubation with the primary mouse monoclonal anti-MDC1 antibody (#M2444, Sigma Aldrich) diluted 1:500 in staining buffer was carried out for 2 h in RT. Slides were then washed four times for 5 min with 0.2% (v/v) PBS-Tween-20 and then incubated with Goat anti-Mouse IgG (H+L) Cross-Absorbed Secondary Antibody, Alexa Fluor 488 (RRID: AB_2534088, #A-11029, Thermo Fisher Scientific) diluted 1:2000 in staining buffer for 1 h at RT. Slides were washed three times for 5 min with 0.2% PBS-Tween- 20, once with PBS and then mounted with Duolink *In Situ* mounting medium with DAPI (#DUO82040, Sigma Aldrich). The same procedure was used for analysis of MDC1 recruitment to irradiation-induced damage sites in RPE-1 cells. To detect the co-localization of MDC1 with γH2AX 2 h-post irradiation, slides were incubated for 1 h in RT with mouse monoclonal Anti-phospho-Histone H2AX (Ser139), clone JBW301 (#05-636, Millipore) primary antibody diluted 1:1000 in staining buffer, or with rabbit polyclonal anti-53BP1 (#ab21083, Abcam) antibody diluted 1:1000 in staining buffer. Slides were washed four times for 5 min with 0.2% PBS-Tween-20 and then incubated with Goat anti-Rabbit IgG (H+L) Cross-Adsorbed Secondary Antibody, Texas Red-X (RRID: AB_2556779, #T-6391, Thermo Fisher Scientific) diluted 1:2000 in staining buffer for 1 h at RT. Z-stack fluorescent images were acquired using the DeltaVision Elite widefield microscope (GE Healthcare Life Sciences). Multiple fields of view were imaged per sample with Olympus 100X/1.40, UPLS Apo, UIS2, 1- U2B836 or Olympus 60X/1.42, Plan Apo N, UIS2, 1-U2B933 objectives and sCMOS camera at the resolution 2048 x 2048 pixels. Deconvolution of the acquired images was performed by the softWoRx DeltaVision software. Images were analyzed and foci quantification analysis was performed using Fiji image processing package of ImageJ (1.52e). Briefly, all nuclei were detected by the “analyze particles” command and all the foci per nucleus were counted with the “find maxima” command. Data were plotted in GraphPad Prism 6 software. For the DNA damage tolerance experiment involving γH2AX detection upon treatment with mitomycin (MMC), olaparib or cisplatin, KB2P1.21 cells were seeded on glass coverslips in a 24-well plate two days before the treatment. To induce replication stress and DNA damage, DMSO, 300 nM MMC, 1 µM olaparib or 100 nM cisplatin were added to the medium for 24 h. Cells were then washed with PBS, fixed with 4% (v/v) PFA/PBS and further processed as described above. Z-stack fluorescent images were acquired using the DeltaVision Elite widefield microscope (GE Healthcare Life Sciences). Multiple fields of view were imaged per sample with Olympus 100X/1.40, UPLS Apo, UIS2, 1-U2B836 objective and sCMOS camera at the resolution 2048 x 2048 pixels. Deconvolution of the acquired images was performed by the softWoRx DeltaVision software. Images were analyzed and foci quantification analysis was performed as described above.

#### Analysis of micronuclei formation

Cells were seeded on coverslips in 24-well plates and treated with DMSO or indicated concentrations of olaparib 24 h later. After 48 h of treatment, cells were washed with PBS and fixed with 4% (v/v) PFA/PBS for 20 min in RT. Cells were then washed 3 times in 0.2% (v/v) PBS-Tween-20 and permeabilized for 20min in 0.2% (v/v) Triton X-100/PBS. Then, slides were washed 3 times with PBS, counterstained with DAPI (1:50000 dilution, #D1306, Life Technologies) and washed 5 times more with PBS before mounting in Fluorescence mounting medium (#S3023, Dako). Z-stack images were acquired using the DeltaVision Elite widefield microscope (GE Healthcare Life Sciences). Multiple fields of view were imaged per sample with Olympus 100X/1.40, UPLS Apo, UIS2, 1-U2B836 objective and sCMOS camera. Frequency of micronuclei positive cells was analyzed manually in Fiji.

#### Total replication speed analysis using EdU incorporation

The total replication speed was assessed using the Click-iT EdU Alexa Fluor 488 Flow Cytometry Assay Kit (#C10420, Thermo Fisher Scientific) and the Click-iT EdU Cell Proliferation Kit for Imaging, Alexa Fluor 488 (#C10337, Thermo Fisher Scientific). For the quantitative flow cytometry-based analysis of EdU incorporation, KB2P1.21 cells were seeded in a 6-well plate 24 h before the experiment. The nascent DNA labeling was carried out by adding 10 µM EdU together with DMSO or 1 µM ATR inhibitor AZ20 in the medium for 30 min. Cells were then washed 3 times with PBS, collected and fixed with 4% (v/v) PFA/PBS. The next steps were carried out according to the manufacturer’s protocol. EdU intensity was measured with the BD FACSCanto II cytometer. FlowJo v10.6.1 software was used to isolate the population of single cells while excluding cell debris and cell doublets, and to export the EdU intensity values for further analysis. EdU intensity values from at least 500 cells/sample in each biological replicate were randomly selected in Excel and plotted in GraphPad Prism 6. Statistical significance was calculated using Mann-Whitney test; ****p<0.0001. For imaging-based analysis of EdU incorporation, the Click-iT EdU Cell Proliferation Kit for Imaging, Alexa Fluor 488 was used. KB2P1.21 cells were seeded on coverslips in a 24-well plate 48 h before the experiment. The pulse labeling was performed as described above. After EdU labeling, cells were washed 3 times with PBS and fixed with 4% (v/v) PFA/PBS. The next steps were carried out according to the manufacturer’s protocol. The images were acquired using the DeltaVision Elite widefield microscope (GE Healthcare Life Sciences). Multiple fields of view were imaged per sample with Olympus 60X/1.42, Plan Apo N, UIS2, 1-U2B933 objective and sCMOS camera. The analysis of EdU intensity was performed using Fiji image processing package of ImageJ (1.52e). Briefly, all nuclei were detected by the “analyze particles” command and EdU incorporation was quantified by measuring the total signal intensity in the nuclear area. Data were plotted in GraphPad Prism 6 software. Representative cells were selected do demonstrate the differences in EdU intensities between the samples.

#### Single molecule DNA fiber assay

Fork progression was measured as described previously in Schmid et al., 2018 with a few modifications. Briefly, asynchronously growing subconfluent KB2P1.21, KB1P-G3 or RPE-1 cells were labeled with 30 μM thymidine analogue 5-chloro-2’-deoxyuridine (CIdU) (#C6891, Sigma-Aldrich) for 20 min, washed three times with warm PBS and subsequently exposed to 250 μM of 5-iodo-2′-deoxyuridine (IdU) for 20 min. In the experiment assessing replication fork stability, IdU pulse was followed by adding medium containing 8 mM HU for 6 h. All cells were then collected by standard trypsinization and resuspended in cold PBS at 3.5 x 10^5^ cells/ml. The labeled cells were mixed 1:5 with unlabeled cells resuspended in cold PBS in the concentration 2.5 x 10^5^ cells/ml. 2.5 μl of this cell suspension were then mixed with 7.5 μL of lysis buffer (200 mM Tris-HCl, pH 7.4, 50 mM EDTA, and 0.5% (v/v) SDS) on a positively-charged microscope slide. After 9 min of incubation at RT, the slides were tilted at an approximately 30-45° angle to stretch the DNA fibers onto the slide. The resulting DNA spreads were air-dried, fixed in 3:1 methanol/acetic acid, and stored at 4 °C overnight. Next day, the DNA fibers were denatured by incubation in 2.5 M HCl for 1 h at RT, washed five times with PBS and blocked with 2% (w/v) BSA in 0.1% (v/v) PBST (PBS and Tween 20) for 40 min at RT while gently shaking. The newly replicated CldU and IdU tracks were stained for 2.5 h at RT using two different anti-BrdU antibodies recognizing CldU (#ab6326, Abcam) and IdU (#347580, Becton Dickinson), respectively. After washing five times with PBST (PBS and Tween 20) the slides were stained with goat the anti-mouse IgG (H+L) Cross-Adsorbed Secondary Antibody, Alexa Fluor 488 (RRID: AB_2534088, #A-11029, Thermo Fisher Scientific) diluted 1:300 in blocking buffer and with the Cy3 AffiniPure F(ab’)₂ Fragment Donkey Anti-Rat IgG (H+L) antibody (#712-165-513, Jackson ImmunoResearch) diluted 1:150 in blocking buffer. Incubation with secondary antibodies was carried out for 1 h at RT in the dark. The slides were washed five times for 3 min in PBST, air-dried and mounted in Fluorescence mounting medium (#S3023, Dako). Fluorescent images were acquired using the DeltaVision Elite widefield microscope (GE Healthcare Life Sciences). Multiple fields of view from at least two slides (technical replicates) of each sample were imaged using the Olympus 60X/1.42, Plan Apo N, UIS2, 1-U2B933 objective and sCMOS camera at the resolution 2048 x 2048 pixels. To assess fork progression CldU + IdU track lengths of at least 120 fibers per sample were measured using the line tool in ImageJ software. Statistical analysis was carried out using GraphPad Prism 6. Replication fork stability was analyzed by measuring the track lengths of CldU and IdU separately and by calculating IdU/CldU ratio. In the replication fork restart experiment stalling with 2 mM hydroxyurea (HU) was performed for 60 min between the CldU and IdU pulse labels. After three washes with warm PBS, IdU pulse was carried out for 40 or 80 min. Samples were then processed as described above. Replication fork restart efficiency was then analyzed by manual counting of CldU tracks only (stalled forks) and CldU + IdU tracks (restarted forks) in ImageJ.

#### Proximity ligation assay (PLA)

KB2P1.21 cells were seeded on sterile glass coverslips in a 24-well plate 48 h before the experiment. EdU labeling was performed by adding 10 μM EdU in the medium for 30 min. Cells were then washed three times with PBS, pre-extracted for 5 min with CSK buffer (25 mM HEPES, pH 7.5, 50 mM NaCl, 1 mM EDTA, 3 mM MgCl2, 300 mM sucrose and 0.5% (v/v) Triton X-100) on ice and fixed with 4% (v/v) PFA/PBS for 15 min at RT. After three washes with PBS, permeabilization was performed with 100% ice cold methanol for 20 min at -20 °C. Slides were then washed three times with PBS and EdU was detected according to the manufacturer’s protocol (Click-iT EdU Cell Proliferation Kit for Imaging, Alexa Fluor 488, #C10337, Thermo Fisher Scientific). Next, slides were blocked with staining buffer (PBS, BSA (2% w/v), glycine (0.15% w/v), Triton X-100 (0.1% v/v)) for 1h at RT. Incubation with the primary antibodies mouse monoclonal anti-MDC1 antibody (#M2444, Sigma Aldrich) diluted 1:500 in staining buffer and rabbit monoclonal anti-PCNA (D3H8P) XP antibody (#13110, Cell Signaling Technology) diluted 1:800 in staining buffer was carried out at 4 °C overnight. The next day, the slides were washed four times with 0.2% (v/v) Tween-20/PBS for 5 min and in situ proximity ligation was performed according to the Duolink Detection Kit protocol (#DUO92101, Sigma Aldrich). Z-stack images were acquired using the DeltaVision Elite widefield microscope (GE Healthcare Life Sciences). Multiple fields of view were imaged per sample with Olympus 100X/1.40, UPLS Apo, UIS2, 1-U2B836 objective and sCMOS camera. All nuclei were detected by the “analyze particles” command, only the EdU positive nuclei were then selected for further analysis. The PLA foci were then quantified with the “find maxima” command. Data were plotted in GraphPad Prism 6.

#### In situ analysis of protein interactions at DNA replication forks (SIRF)

RPE-1 cells were seeded on sterile glass coverslips in a 24-well plate 48 h before the experiment in the density of 30 000 cells/well. EdU labeling was performed by adding 25 μM EdU in the medium for 10 min. After three washes with PBS, replication stress was induced in the HU-treated samples by adding 2 mM HU for 2 h. Cells were then washed twice with PBS and nuclei were pre-extracted with CSK buffer (10 mM PIPES pH 7, 0.1 M NaCl, 0.3 M sucrose, 3 mM MgCl2, 0.5 % (v/v) Triton X-100) on ice for 5 min. After pre-extraction, cells were washed with PBS and fixed in 4% (v/v) PFA/PBS for 15 min at RT. After three washes with PBS, permeabilization was performed with 0.2 % (v/v) Triton X-100 for 15 min at RT. Then, click reaction was performed by adding the click reaction buffer (100 mM Tris pH 8, 4 mM CuSO4, 100 mM sodium ascorbate, 50 µM biotin-azide) to the samples and incubating at 37 °C for 2 h. Slides were then incubated in the blocking buffer (PBS, BSA (2% w/v), glycine (0.15% w/v), Triton X-100 (0.1% v/v)) for 1 h at 37 °C, followed by incubation with primary antibodies mouse anti-MDC1 (#M2444, Sigma Aldrich, diluted 1:200) and rabbit anti-biotin (D5A7, #5597, Cell Signaling, diluted 1:1000) overnight at 4 °C. The next steps are identical to the steps described in the PLA protocol above.

#### Immunoblotting

200.000 RPE-1 cells were seeded in a 6-well plate 24 h prior the experiment. Replication stalling was induced by adding 1 mM hydroxyurea (HU) into the medium for 24 h. Cells were then washed with PBS, trypsinized and collected in the 15 ml tubes. After washing with PBS, cells were lysed for 40 min in RIPA buffer supplemented with Halt Protease and Phosphatase Inhibitor Cocktail (100x) (#78420, Thermo Fisher Scientific) while briefly vortexed every 10 min. Lysates were then centrifuged at 10.000 rpm for 10 min at 4 °C and the supernatant was collected to determine protein concentration using Pierce BCA Protein Assay Kit (#23225, Thermo Fisher Scientific). Before loading, protein lysates were denatured at 95 °C for 5 min in 6x SDS sample buffer. Proteins were separated by SDS/PAGE in 10% gel before wet transfer to 0.45 µm nitrocellulose membranes (GE Healthcare) and blocked in 5% dry milk powder in TBS-T (100 mM Tris, pH 7.5, 0.9% NaCl, 0.05% Tween-20). Membranes were incubated with the following primary antibodies: rabbit polyclonal anti-RPA32 (1:1000, #A300-244A, Bethyl Laboratories), rabbit recombinant monoclonal anti-phospho RPA32 (S4/S8) (1:1000, #A700-009, Bethyl Laboratories), rabbit polyclonal anti-CHK1 (1:5000, #NB100-464, Novus Biologicals), rabbit monoclonal anti-phospho CHK1 (S345) (133D3) (1:1000, #2348, Cell Signaling), rabbit polyclonal anti-KAP1 (1:1000, #NB500-158, Novus Biologicals), rabbit polyclonal anti-phospho KAP1 (S824) (1:1000, #A300-767A, Bethyl Laboratories), rabbit polyclonal anti-γ-Tubulin (1:1000, #5886, Cell Signaling) and mouse monoclonal anti-PARP1 (1:500, #MA3-950, Thermo Fisher Scientific). The antibodies were diluted in 5% milk in TBS-T and incubated with membranes for 2 h at RT. After three 5 min washes in TBS-T, anti-mouse or anti-rabbit Horseradish Peroxidase (*HRP*)-linked secondary antibodies (Cell Signaling, dilution 1: 2500) were applied for 1 h at room temperature. Images were acquired using Vilber FUSION FX chemiluminescent imager.

#### Transmission electron microscopy of replication intermediates (RI)

The procedure was performed as described previously by Zellweger and Lopes, 2018 with minor modifications. A total of 2.5–5.0 x 106 asynchronously growing subconfluent RPE-1 cells were harvested by trypsinization and resuspended in 10 mL of cold PBS. DNA was cross-linked by exposing the living cells twice to 4,5′,8-trimethylpsoralen at a final concentration of 10 μg/mL followed by 3 min irradiation pulses with UV 365-nm monochromatic light (UV Stratalinker 1800, Agilent Technologies). The cells were then washed repeatedly with cold PBS and lysed with a cell lysis buffer (1.28 M sucrose, 40 mM Tris-Cl, pH 7.5, 20 mM MgCl2, and 4% (v/v) Triton X-100). The nuclei were then digested in a digestion buffer (800 mM guanidine-HCl, 30 mM Tris-HCl, pH 8.0, 30 mM EDTA, pH 8.0, 5% (v/v) Tween 20, and 0.5% (v/v) Triton X-100) supplemented with 1 mg/mL proteinase K at 50 °C for 2 h. Genomic DNA was extracted with a 24:1 Chloroform:Isoamyl alcohol mixture by phase separation (centrifugation at 8.000 rpm for 20 min at 4 °C) and precipitated by addition of equal amount of isopropanol to the aqueous phase, followed by another centrifugation step (8.000 rpm for 10 min at 4 °C). The obtained DNA pellet was washed once with 1 mL of 70% ethanol, air-dried at RT, and resuspended by overnight incubation in 200 μL TE (Tris-EDTA) buffer at RT. 12 μg of the extracted genomic DNA was digested for 5 h at 37 °C with 100 U restriction enzyme PvuII high-fidelity (#R3151S, New England Biolabs). The digest was cleaned up using a silica bead DNA gel extraction kit (#K0513, Thermo Fisher Scientific). The benzyldimethylalkylammonium chloride (BAC) method was used for native spreading of the DNA on a water surface and then loading it on carbon-coated 400-mesh magnetic nickel grids. After the spreading procedure, the electron density of the DNA was increased by platinum coating with the platinum-carbon rotary shadowing technique using the MED 020 High Vacuum Evaporator (Bal-Tec). The grids were then scanned in a semi-automated fashion using a transmission electron microscope (FEI Thalos 120, LaB6 filament) at high tension ≤ 120 kV and pictures were acquired with a bottom mounted CMOS camera BM-Ceta (4000 x 4000 pixels). The images were processed with MAPS Version 3.14 (Thermo Fisher) and analyzed using MAPS Offline Viewer Version 3.14.11 (Thermo Fisher). Mean + SD values of analyzed frequency of RIs obtained from three independent replicates were plotted and statistics was performed in GraphPad Prism 9 using unpaired t test.

### QUANTIFICATION AND STATISTICAL ANALYSIS

Statistical parameters including sample size, number of biological replicates, applied statistical tests and statistical significance are reported in the figures, corresponding figure legends and Method Details sections.

#### CRISPR/Cas9 genetic screens

Sequence alignment and enrichment analysis done by comparing day 0 vs PARPi-treated population from six replicates (independent mutagenesis) was carried out using MAGeCK software (Li et al., 2014).

#### Clonogenic Assays

See Figures 1B, 1C, S1F, S2B, S2C. All experiments indicated in these figures were performed as at least two individual biological replicates and graphs were drawn from these data using GraphPad prism 6. Cell survival in each condition was normalized to the corresponding DMSO-treated control. Statistical analysis was performed in Graphpad Prism using a Two-way ANOVA followed by Dunnet’s test; *p <0.05, **p <0.01, ***p<0.001; ****p<0.0001. Analysis of the competition assays in Figures S1D and S1H was carried out by comparing the frequency of modifications in gRNA1/2-targeted sites in DNA samples from *Mdc1*-targeting gRNA1/2 cells before and after PARPi treatment, and NT gRNA cells collected at day 0 of the experiment.

#### CTB proliferation assay

See Figure S4B, all the measured values were obtained from at least three independent biological replicates. Fluorescence measured in each condition was normalized to the background fluorescence of CellTiter Blue reagent incubated in an empty well with growth medium. The date were plotted using GraphPad Prism 6.

#### In vivo studies

See Figures 1F and S2E. 5 mice were used in each of the four group (Non-targeting gRNA or *Mdc1*-targeting gRNA1; vehicle- or AZD2461-treated) in the Figure 1F. 10 mice were used in each group (Non-targeting gRNA or *Mdc1*-targeting gRNA1; vehicle- or AZD2461-treated) in the Figure S2E. Kaplan Meyer survival curves were plotted and statistical analysis was performed using log-rank test in GraphPad prism 6.

#### Immunofluorescence

See Figures 2A, 2D, 2F, 2G, S1B, S3E, S4A and S4C. Each condition was stained as indicated in the method details section. GE DeltaVision fluorescent microscope with 60x or 100x objectives with immersion oil were used to acquire images. Each image was taken in at least 6 Z-layers and maximum intensity Z-projection was performed in Fiji to create a single layer for quantification or for preparation of representative images. Foci were quantified in the Figures 2A, 2D, 2F, S1B S3E and S4C using the following procedure: Nuclei were segmented by manual intensity thresholding in the DAPI channel and separation using the “watershed’’ function. Detection of individual nuclei for analyzes was performed by “Analyze particles” function. Number of foci per cell was quantified by “find maxima” function in each defined particle. A minimum of 100 cells were quantified from each condition and the average number of foci were plotted in GraphPad Prism 6. Statistical analysis in the Figures 2D, S3E and S4C was performed using Mann-Whitney test; ****p<0.0001. Analyzes in the Figure 2A was performed using Two-way ANOVA followed by Tukey’s multiple comparison test; *p <0.05. Analyzes in the Figure 2F was performed using Two-way ANOVA followed by Dunnet’s multiple comparison test; ns = non-significant, *p <0.05, **p <0.01. In Figure 2G, micronuclei positive cells were counted manually and statistical significance was calculated using the Two-way ANOVA test; *p <0.05, **p <0.01, ***p<0.001; ****p<0.0001.

#### Single molecule DNA fiber assay

See Figures 2B, 3C, 3D, 4A, 4B, 4C, 4D, 4E, 4F, S3A, S3B, S3C and S3D. The nascent DNA pulse labeling, fiber spreading and processing of slides was performed as described in Method details section. Minimum of 100 tracks were measured from two microscopy slides (technical replicates) in each biological replicate. In Figure 3D, 90-170 individual track lengths were measured from two independent biological replicates. Track lengths were plotted in GraphPad Prism 6 and statistical significance was calculated using Mann-Whitney test; *p <0.05, **p <0.01, ***p<0.001; ****p<0.0001.

#### Immunoblotting

See Figures 2E and S4D. Immunoblotting experiments were performed at least twice. Representative images are shown.

## ADDITIONAL RESOURCES

There are no additional resources used in this manuscript.

---

**Figure S1.**
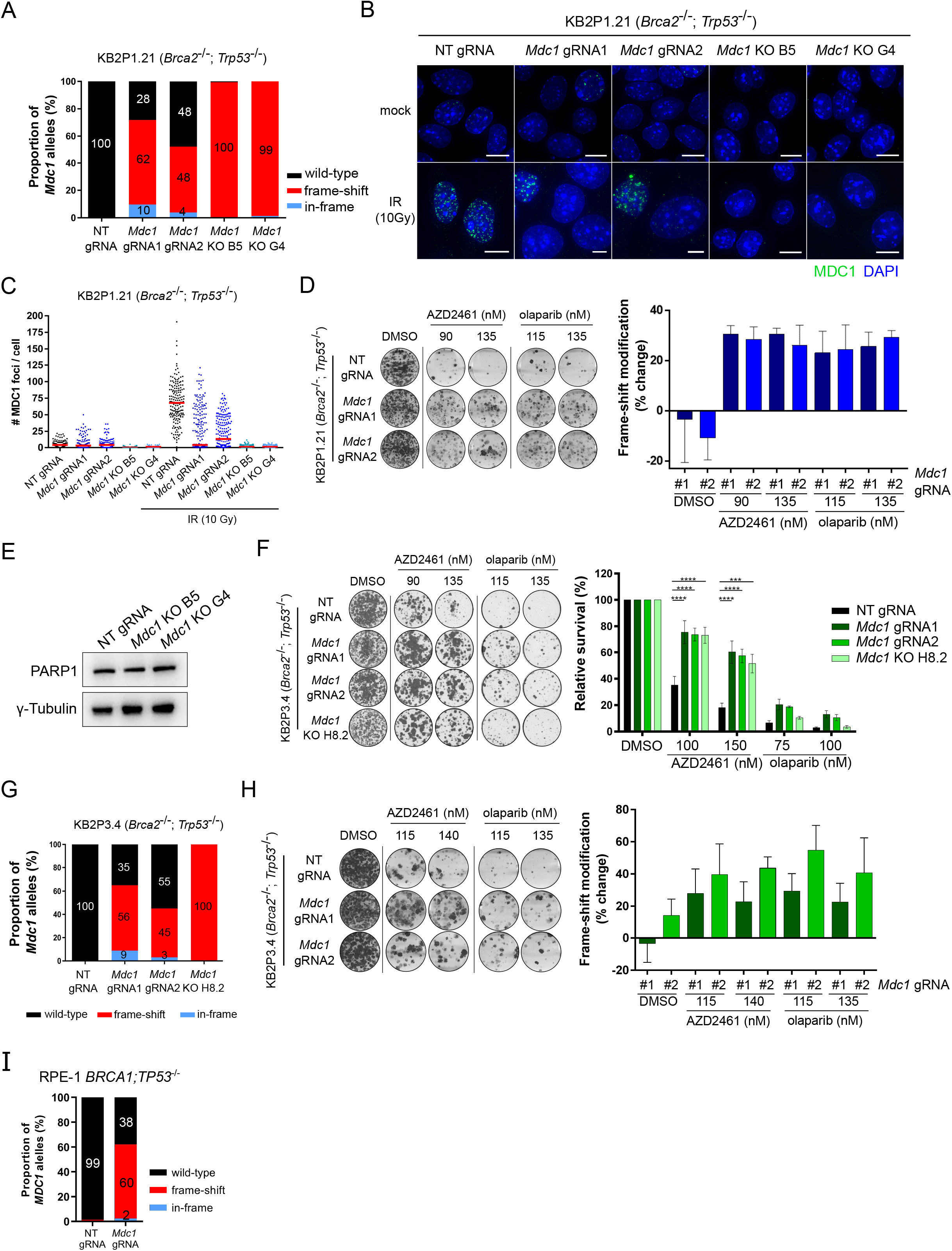
MDC1 deficiency promotes PARPi resistance *in vitro*. Related to Figure 1. **A)** TIDE analysis showing the modification rate of the *Mdc1* gene in the KB2P1.21 cells expressing non-targeting (NT) or *Mdc1*-targeting gRNA. **B)** Immunofluorescence of MDC1 (green) in the presence or absence of irradiation. Scale bars represent 10 µm. **C)** Quantification of MDC1 foci formation in absence or presence of irradiation in KB2P1.21 cells. Median of measured values is shown. **D)** Competition assay in control and *Mdc1*-targeted polyclonal cell lines with two PARP inhibitors. Representative images (left) and TIDE analysis of relative change in frequency of frame-shift modifications in *Mdc1* gene compared to the KB2P1.21 cells expressing NT gRNA (right). Data represent Mean ± SD of three biological replicates. **E)** Western blotting of PARP1 and γ-Tubulin in NT gRNA and *Mdc1* KO B5 and G4 cells. Similar results were observed in two independent experiments. **F)** Clonogenic survival assay with representative images (left) and quantification of cell survival (right). Mean + SEM of three independent experiments is shown. **G)** TIDE analysis demonstrating the *Mdc1*-targeting efficiency in KB2P3.4 cells. **H)** Competition assay in polyclonal KB2P3.4 cell lines with representative images (left) and quantification of change in the frequency of frame-shift modifications following the treatment with PARPi (right). Mean ± SD of three independent experiments is shown. **I)** TIDE analysis showing the modification rate of the *MDC1* gene in the RPE-1 cells expressing non-targeting (NT) or *MDC1*-targeting gRNA.

**Figure S2.**
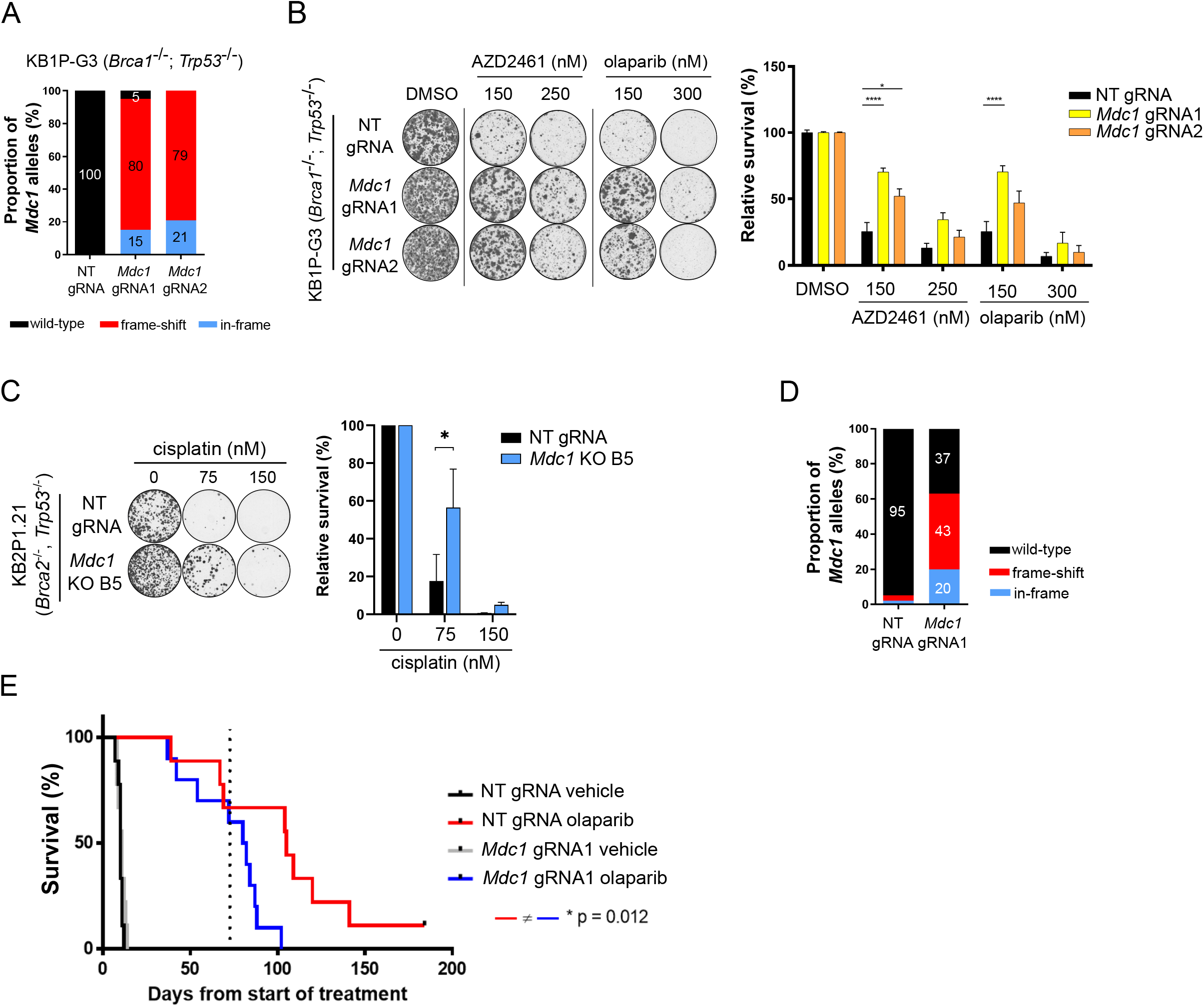
MDC1 deficiency promotes chemoresistance *in vitro* and *in vivo*. Related to Figure 1. **A)** TIDE analysis showing the frequency of WT and modified *Mdc1* alleles in KB1P-G3 cells. **B)** Clonogenic survival assay of KB1P-G3 cells expressing NT, or *Mdc1*-targeting gRNAs upon treatment with PARP inhibitors. Representative images (left) and quantification (right) are shown. Mean ± SEM of at least three independent experiments is shown. **C)** Clonogenic survival assay with representative images (left) and quantification of cell survival (right). Mean + SD of at least two independent experiments is shown. **D)** TIDE analysis showing the rate of modifications in *Mdc1* gene in tumors derived from the KB2P 3D organoids. **E)** Kaplan-Meier curve of overall survival of mice treated with vehicle or PARPi olaparib.

**Figure S3.**
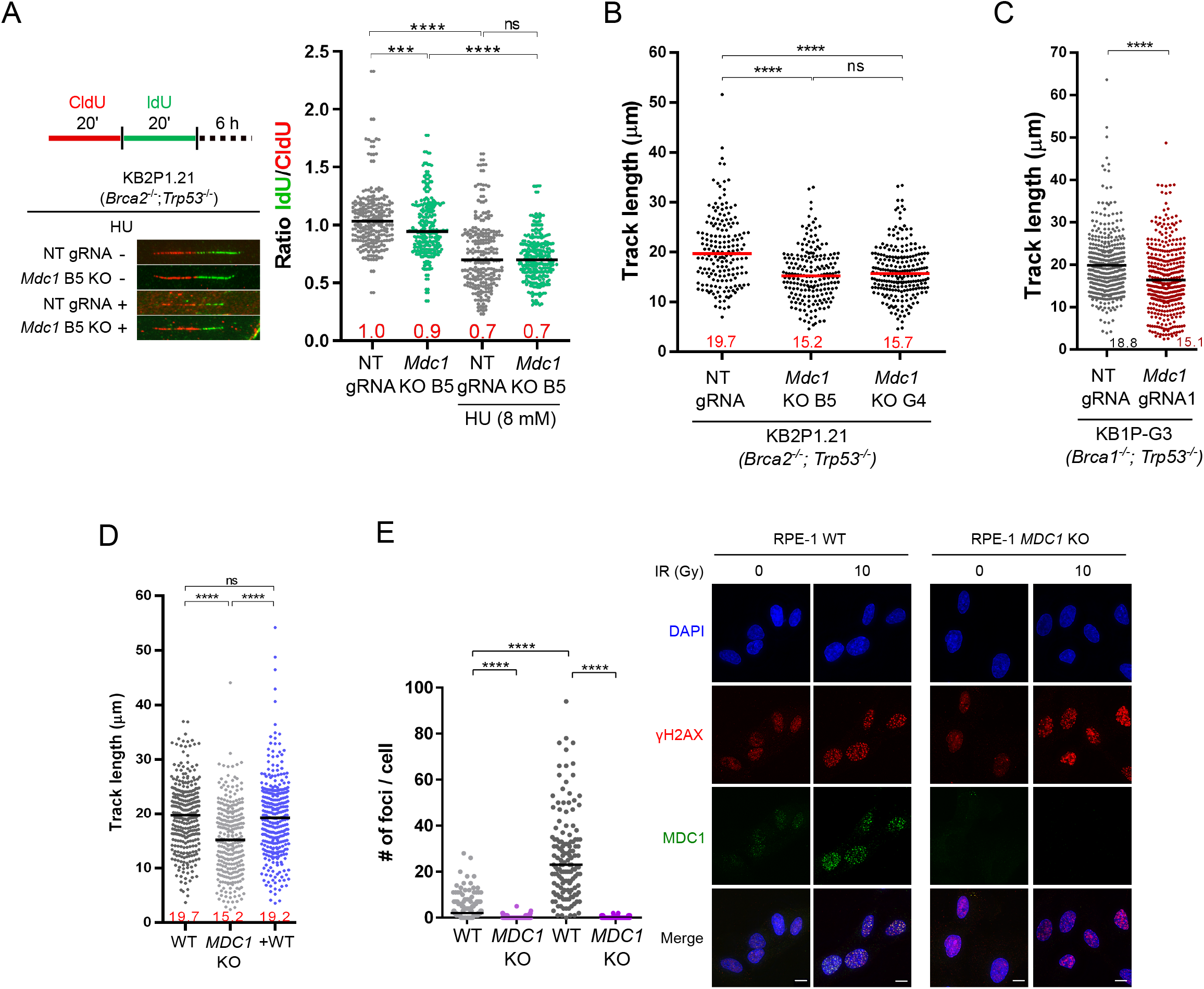
MDC1 deficiency does not restore stability of replication forks, but reduces fork progression. Related to Figure 2. **A)** DNA fiber assay showing RF degradation in KB2P1.21 cells upon HU-induced RF stalling. Minimum of 120 forks per sample were analyzed and the IdU/CldU ration was calculated. Median of values measured from two independent experiments is shown. **B)** DNA fiber assay showing reduced RF speed in KB2P1.21 *Mdc1* knockout lines B5 and G4. Track lengths of at least 100 forks were measured. Median of track lengths is shown. Similar results were observed in at least three biological replicates. **C)** Track lengths analysis in KB1P-G3 cells expressing NT, or *Mdc1*-targeting gRNA. Minimum of 100 forks per sample were analyzed and median of values measured from two independent experiments is shown. **D)** DNA fiber assay in *MDC1* WT, *MDC1* KO and GFP-tagged MDC1 WT complemented RPE-1 cells. The median of track lengths from two independent experiments is shown. **E)** Immunofluorescence analysis showing the loss of MDC1 IRIF formation in RPE-1 *MDC1* knockout cell line. The scale bar represents 10 µm.

**Figure S4.**
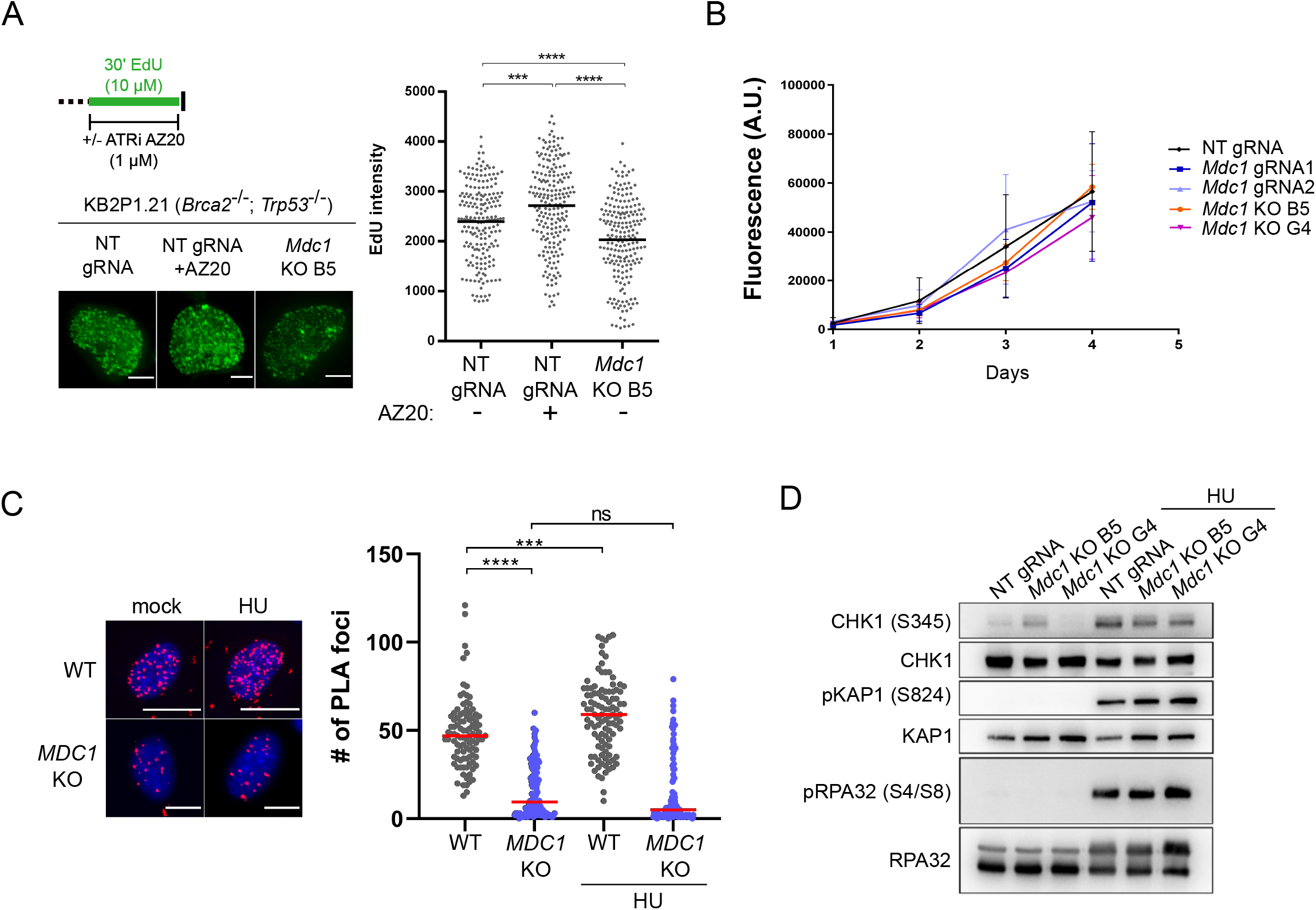
MDC1 associates with active replication forks and regulates their speed independently of new origin firing and changes in cell proliferation rate. Related to Figure 2. **A)** Imaging-based analysis of EdU incorporation upon release of new origins by ATR inhibitor AZ20 or upon *Mdc1* KO in KB2P1.21 cells. The scale bars represent 5 µm. **B)** Proliferation rate of KB2P1.21 cells expressing NT gRNA, two *Mdc1*-targeting gRNAs and two KO cell lines B5 and G4. Mean ± SD of at least three biological replicates is shown. **C)** Analysis of MDC1 localization at active replication forks by SIRF. Representative images (left) and quantification (right) are shown. Similar results were obtain from at least two independent experiments. Scale bars represent 5 µm. **D)** Western blotting of DDR markers before and after 1 mM HU for 24 h in NT gRNA, B5 and G4 KB2P1.21 cells. Similar results were obtained from two independent experiments.

**Table S1.** List of the ranked highest-scoring gene candidates identified in 6 biological replicates of the CRISPR/Cas9-based PARPi resistance screens.

| id | num | neg\|score | neg\|p-value | neg\|fdr | neg\|rank | neg\|goodsgrna | pos\|score | pos\|p-value | pos\|fdr | pos\|rank | pos\|goodsgrna |
| --- | --- | --- | --- | --- | --- | --- | --- | --- | --- | --- | --- |
| Ankmy2 | 3 | 0.99862 | 0.9986 | 0.999895 | 21026 | 0 | 1.0883e-07 | 2.3509e-07 | 0.00495 | 1 | 3 |
| Mdc1 | 3 | 0.94771 | 0.94754 | 0.999895 | 19773 | 0 | 5.7779e-07 | 2.1158e-06 | 0.022277 | 2 | 2 |
| Ric8 | 3 | 0.99998 | 0.99998 | 0.999978 | 21058 | 0 | 2.1488e-05 | 5.4305e-05 | 0.352723 | 3 | 3 |
| Usp51 | 3 | 0.9022 | 0.90183 | 0.999895 | 18836 | 0 | 2.5589e-05 | 6.7e-05 | 0.352723 | 4 | 2 |
| H2afx | 3 | 0.99996 | 0.99996 | 0.999978 | 21057 | 0 | 3.6542e-05 | 9.8032e-05 | 0.412871 | 5 | 3 |
| Ube3c | 3 | 0.99995 | 0.99995 | 0.999978 | 21056 | 0 | 4.6271e-05 | 0.00012812 | 0.44967 | 6 | 3 |
| Gm20854 | 3 | 0.99994 | 0.99994 | 0.999978 | 21055 | 0 | 5.9992e-05 | 0.00017702 | 0.532532 | 7 | 3 |
| Cdrt4 | 3 | 0.59565 | 0.68782 | 0.999895 | 14294 | 1 | 7.4532e-05 | 0.00021793 | 0.573639 | 8 | 2 |
| Lcn5 | 3 | 0.70296 | 0.7458 | 0.999895 | 15551 | 1 | 0.00010477 | 0.00031902 | 0.746425 | 9 | 2 |
| Sprr1b | 3 | 0.81303 | 0.8202 | 0.999895 | 17072 | 1 | 0.00012422 | 0.00037826 | 0.796535 | 10 | 2 |
| Brk1 | 3 | 0.9909 | 0.99088 | 0.999895 | 20825 | 0 | 0.00014143 | 0.00042245 | 0.808731 | 11 | 3 |
| Olfr1276 | 3 | 0.84304 | 0.8449 | 0.999895 | 17610 | 1 | 0.0001739 | 0.00051555 | 0.827905 | 12 | 1 |
| Tmem170 | 3 | 0.99323 | 0.99319 | 0.999895 | 20869 | 0 | 0.00019774 | 0.00058184 | 0.827905 | 13 | 3 |
| Mrgprb2 | 3 | 0.070688 | 0.15692 | 0.979018 | 3364 | 1 | 0.00022359 | 0.0006646 | 0.827905 | 14 | 2 |
| 3830403N18Rik | 2 | 0.99977 | 0.99974 | 0.999933 | 21054 | 0 | 0.00023346 | 0.00068716 | 0.827905 | 15 | 2 |
| Wbscr27 | 3 | 0.99973 | 0.99971 | 0.999933 | 21053 | 0 | 0.00026881 | 0.00077932 | 0.827905 | 16 | 3 |
| Palb2 | 3 | 0.88625 | 0.88582 | 0.999895 | 18493 | 0 | 0.00027327 | 0.00079107 | 0.827905 | 17 | 1 |
| Trpc5 | 3 | 0.98871 | 0.98872 | 0.999895 | 20760 | 0 | 0.00028127 | 0.00080565 | 0.827905 | 18 | 3 |
| Vmn1r60 | 3 | 0.99967 | 0.99964 | 0.99993 | 21052 | 0 | 0.00032748 | 0.00092836 | 0.827905 | 19 | 3 |
| Clec4e | 3 | 0.99782 | 0.99779 | 0.999895 | 21002 | 0 | 0.00033109 | 0.00093824 | 0.827905 | 20 | 3 |

## Notes

### Competing Interest Statement

The authors have declared no competing interest.

